# Morphology and gene expression profiling provide complementary information for mapping cell state

**DOI:** 10.1101/2021.10.21.465335

**Authors:** Gregory P. Way, Ted Natoli, Adeniyi Adeboye, Lev Litichevskiy, Andrew Yang, Xiaodong Lu, Juan C. Caicedo, Beth A. Cimini, Kyle Karhohs, David J. Logan, Mohammad H. Rohban, Maria Kost-Alimova, Kate Hartland, Michael Bornholdt, Srinivas Niranj Chandrasekaran, Marzieh Haghighi, Erin Weisbart, Shantanu Singh, Aravind Subramanian, Anne E. Carpenter

## Abstract

Morphological and gene expression profiling can cost-effectively capture thousands of features in thousands of samples across perturbations by disease, mutation, or drug treatments, but it is unclear to what extent the two modalities capture overlapping versus complementary information. Here, using both the L1000 and Cell Painting assays to profile gene expression and cell morphology, respectively, we perturb A549 lung cancer cells with 1,327 small molecules from the Drug Repurposing Hub across six doses, providing a data resource including dose-response data from both assays. The two assays capture both shared and complementary information for mapping cell state. Cell Painting profiles from compound perturbations are more reproducible and show more diversity, but measure fewer distinct groups of features. Applying unsupervised and supervised methods to predict compound mechanisms of action (MOA) and gene targets, we find that the two assays provide a partially shared, but also a complementary view of drug mechanisms. Given the numerous applications of profiling in biology, our analyses provide guidance for planning experiments that profile cells for detecting distinct cell types, disease phenotypes, and response to chemical or genetic perturbations.

## Introduction

In a profiling experiment, biologists measure high-dimensional readouts from biological samples (e.g. single cells, organoids, tissue, whole organisms). The resulting profile contains measurements of hundreds to thousands of individual features that together form a systems biology representation of the sample of interest. Automation now allows biologists to probe thousands of chemical and genetic perturbations to assess their phenotypic impact (Dixit et al., 2016; Keenan et al., 2018; Subramanian et al., 2017a). Therefore, perturbational profiling results in a large number of samples measured across a common set of high-dimensional features. Biologists can then apply data mining and machine learning to these datasets to detect and quantify the similarities and differences among samples. These approaches have the potential to advance drug discovery, functional genomics, and precision medicine, for example, by annotating uncharacterized small molecules, cataloging the mechanistic outcome of gene editing, and testing the impact of specific perturbations on disease-associated phenotypes (Chandrasekaran et al., 2021; Malone et al., 2020; Musa et al., 2018).

Biologists can access different aspects of cell state through multiple profiling assays that capture different biological landscapes: DNA, RNA, epigenetic marks, metabolites, microbiota, proteins, kinases, and spatial information (Cazaly et al., 2019; Di Minno et al., 2021; Litichevskiy et al., 2018; Ottestad et al., 2020; Wang and Ma, 2015; Waylen et al., 2020; Yang et al., 2020). Some profiling approaches measure multiple data modalities in the same assay and are dubbed multi-omic or multi-modal assays (Cao et al., 2018; Hu et al., 2018); others pool and de-multiplex perturbations to increase throughput (McFarland et al., 2020); still others test a single readout (e.g. viability) but across hundreds of cell types to yield a profile (Garnett et al., 2012; Yu et al., 2016).

Gene expression and cell morphology are currently the two highest-throughput, lowest cost, high-dimensional profiling data types for mammalian cells (Bray et al., 2016; Gustafsdottir et al., 2013; Subramanian et al., 2017a). These readouts measure fundamentally different aspects of biology. In the L1000 bead-based assay, probes targeting 978 genes measure mRNA transcript levels (gene expression) in a cell population (Subramanian et al., 2017a). In the Cell Painting assay, after treating cells with six fluorescent dyes to mark eight cellular compartments (**Figure 1A**), biologists use a microscope to image five channels and use software to analyze and extract several thousand morphology measurements from each cell (Bray et al., 2016). Both the gene expression and morphology landscapes change as cells respond to perturbations.

**Figure 1.**
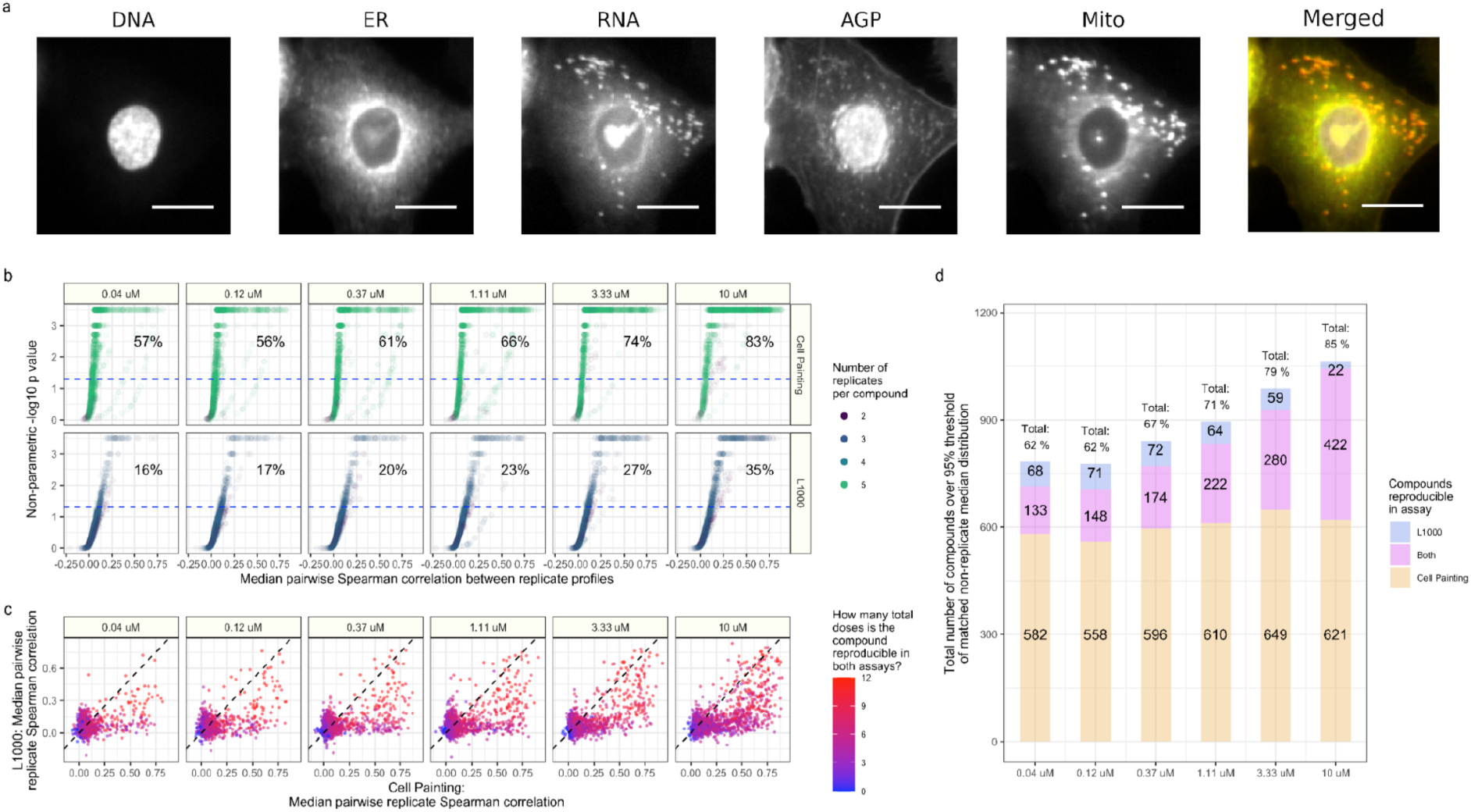
Cell Painting and L1000 data provide complementary measurements of compound perturbations across doses. **(a)** An example Cell Painting image of a single A549 lung cancer cell measured across five channels. We show a merged representation as well. ER = endoplasmic reticulum; Mito = mitochondria; AGP = actin, Golgi, plasma membrane. Scale bar is 20 μm. **(b)** The percent replicating metric measures the percentage of profiles that correlate with each other to a level higher than a carefully-matched and randomly-sampled null distribution. See methods for full details about sampling and data processing. The dotted blue line indicates the 95th percentile of the matched non-replicate distribution. **(c)** Median pairwise replicate Spearman correlations between profiles measured by the L1000 assay (y axis) and Cell Painting assay (x axis). The dotted black line is the line y = x, so anything above is measured with a higher replicate correlation in L1000 and vice versa. **(d)** The L1000 and Cell Painting assays reproducibly measure a complementary set of compound perturbations. The three numbers represent (from top to bottom) the number of compounds unique to L1000, the number of compounds captured in both assays, and the number of compounds unique to Cell Painting that have median pairwise replicate correlations above the randomized non-replicate correlation threshold.

Scientists have used individual profiling modalities to advance a variety of drug discovery applications, including improving screening library diversity, predicting cytotoxicity, prioritizing compounds for follow-up study, and inferring the mechanism of action of chemicals (Feng et al., 2009; Filzen et al., 2017; Lapins and Spjuth, 2019; Ljosa et al., 2013; Nyffeler et al., 2020; Perlman et al., 2004; Wawer et al., 2014; Way et al., 2021b). Integrating gene expression and morphology profiles with chemical structures revealed that each data type provides complementary information for predicting a drug’s mechanism of action (Haghighi et al., 2021; Nassiri and McCall, 2018), for predicting the effects of perturbations (Caicedo et al., 2021a), and for identifying nuisance compounds that can lead to false hits (Dahlin et al., 2021). As well, to some degree, gene expression and morphology datasets contain sufficient information to predict changes in each other (Haghighi et al., 2021; Nassiri and McCall, 2018; Wakui et al., 2022).

However, the field lacks a systematic study evaluating both assays’ information content in terms of distinct versus overlapping signals, diversity of cell states, and performance in useful tasks. The ability of a profiling assay to accomplish a biological task is a function of its technical reproducibility, its inherent information content, properties of bioinformatics pipelines, and natural biological variation. Therefore, our goal in this study is to determine how the assays compare with each other on useful biological tasks given all those sources of variation/noise and current best practices in data processing.

In this study, we collected L1000 and Cell Painting readouts from a common set of 1,327 different Drug Repurposing Hub compounds and controls across six doses representing 511 different mechanism-of-action (MOA) classes and 720 different gene targets (Corsello et al., 2017). After data processing, we observed that while Cell Painting suffers from more batch and well position effects that must be carefully adjusted, the assay showed higher profile reproducibility than L1000. While L1000 includes more independent feature groups than Cell Painting, the latter provides a higher sample diversity. We test the practical implications of these properties by predicting compound MOA and gene targets using two approaches: an unsupervised matching approach and supervised deep learning in which we train top-performing models from a recent Kaggle competition (Kaggle.com et al., 2020). Both assays predict a small set of mechanisms consistently well and certain mechanisms are better captured in one assay or the other. MOA prediction is a challenging task where even small improvements could impact drug discovery. In summary, we find that Cell Painting and L1000 each reproducibly measure a partly overlapping, partly distinct set of compound mechanisms. Based on this analysis we conclude that measuring both molecular and cellular phenotypes increases the ability to capture relevant biological mechanisms from unbiased compound screens.

## Results

### Measuring and processing morphology and gene expression data

One strategy to interrogate biological processes is to measure cell responses to various perturbations in high-throughput, high-dimensional profiling assays. Profiling assays vary in style and measurement, and the sensitivity and resolution with which they capture important biological signals depends on the assay chosen. In this experiment, we asked whether measuring the same perturbations using fundamentally different kinds of profiling assays provides advantages. We therefore created and analyzed two profiling datasets that capture different types of information: gene expression with the L1000 assay and cell morphology with the Cell Painting microscopy assay (Bray et al., 2016; Gustafsdottir et al., 2013; Subramanian et al., 2017a). We show raw data for both Cell Painting and L1000 assays in **Figures S1 and S2**, respectively.

We perturbed A549 lung cancer cells with a common set of 1,327 compounds and controls from the Drug Repurposing Hub (Corsello et al., 2017). We selected compounds that were in current clinical use or in advanced clinical testing and chose them to represent a diversity of mechanisms based on Drug Repurposing Hub annotations (Corsello et al., 2017). We measured a total of 1,258 compounds across six doses (usually, 0.04 μM to 10 μM); the remainder had three to five doses. We provide compound annotations for MOAs and gene targets in **Table S1**.

We perturbed the cells under consistent experimental conditions, including the same 384-well plate layouts (**Figure S3A**). We exposed cells to compounds for 24 hours prior to L1000 profiling and for 48 hours prior to Cell Painting, using standard assay time points based on past experience. At these time points and doses, we did not observe high amounts of cell death which otherwise might have impeded our ability to acquire cell responses to perturbation (**Figure S4**).

The compounds were arrayed in 25 different plate maps (compound layouts), and, in most cases, we collected three replicate plates per plate map for L1000 and five replicate plates per plate map for Cell Painting, given its lower cost per plate. Each replicate plate contained 56 different compounds in six doses plus 24 dimethyl sulfoxide (DMSO) negative controls and 24 proteasome inhibitor positive controls.

We applied standard data processing pipelines for each assay (see Methods) to normalize and transform the data prior to downstream analyses (**Figure S3B**). Due to the limitations of the compound dispensing equipment, it was unfortunately infeasible to control for plate layout artifacts by scrambling perturbation locations within each plate across replicates. Indeed, we observed plate position effects in the Cell Painting data, particularly in edge wells (**Figure S5**). Therefore, we applied a spherize transform using negative control DMSO wells to combine data across batches and adjust for these plate-position effects. Spherizing, also known as whitening, adjusts all profiles such that the DMSO wells are transformed to have an identity covariance matrix (Ando et al., 2017; Kessy et al., 2018) (see Methods for more details). In all the downstream analyses, we use spherized Cell Painting profiles and the original, unspherized L1000 profiles unless indicated otherwise (L1000 did not benefit from spherizing, see **Figure S5, bottom**).

### Assessing profile reproducibility in L1000 and Cell Painting assays

To study a perturbation’s function, a scientist must reliably and robustly measure its biological effect. Therefore, we introduced and calculated a reproducibility metric, based on median pairwise Spearman correlations, which we call “percent replicating” (**Figure S6**). Specifically, percent replicating captures the percentage of profile replicates (treatment of the same compound measured at the same dose) that are more similar to one another than to a randomly-permuted null distribution that adjusts for dose, sample size, and well position (see Methods for complete details).

As expected, percent replicating increased with dose in both assays, as higher concentrations of drug are more likely to impact cell systems (**Figure 1B**). However, we observed much higher percent replicating scores in Cell Painting (57% to 83%, from lowest to highest dose) compared to L1000 (16% to 35%, from lowest to highest dose) (**Figure 1B**); 35% at the highest dose for L1000 is consistent with prior observations (Subramanian et al., 2017a). We provide median pairwise correlations and percent replicating p values for all compounds per assay in **Table S2**.

We expected to observe higher percent replicating scores for Cell Painting over L1000 because we had, typically, five replicates of Cell Painting and only three replicates of L1000 per treatment as per standard assay guidelines. Indeed, a subsampling experiment that randomly sampled Cell Painting replicates to match the number of L1000 replicates reduced percent replicating (from 57% to 37% and 83% to 67%, from lowest to highest dose), although still higher than L1000 (**Figure S7**). Another possible explanation for higher Cell Painting reproducibility metrics is that plate layout effects artificially increased replicate correlations preferentially for that modality versus L1000. Indeed, we observed that percent replicating increased if we failed to adjust our null distribution for well position, and decreased if we failed to correct for plate-position effects (**Figure S7**). However, reproducibility metrics were robust to edge well filtering and different null distributions, as we observed similar performance if we 1) prefiltered edge wells as quality control, 2) adjusted null distributions only for dose and not sample size, and 3) dropped dose altogether when sampling the null distribution (**Figure S8**). These results underscore the importance of maximizing replicate treatments, proper construction of null distributions, and proper profile normalization in Cell Painting. The same normalization did not improve scores for L1000.

Plate layout effects are a serious concern in profiling experiments where practical reasons require scientists to measure replicates of a sample at the same well position across physical plates; it is known that the location on the plate, especially distance from the edge of the plate, can impact many cell phenotypes. Therefore, to more closely study the impact of plate position on pairwise correlations, we performed a non-replicate diffusion analysis in which we systematically increased the well neighborhood size in calculating the null distribution of non-replicate correlations (see Methods). Briefly, we started with a diffusion size 0, which looks at the non-replicate correlations of different samples that are in the exact same well position, across different plate maps. As we increased diffusion (the well neighborhood size) to include adjacent and nearby wells, we observed a slight dampening of non-replicate correlations (**Figure S9A**). While this analysis revealed increased plate-position effects in Cell Painting compared to L1000, this bias is relatively small compared to the overall replicate correlation signal (**Figure S9B**). Taken together, plate layout effects do impact profiling assays but are unlikely to have driven the signal we observed in this experiment. Nevertheless, when possible, we recommend scrambling replicates across wells to avoid this potential confounding effect.

Comparing median pairwise replicate correlations across individual treatments, we observed that most compounds have higher correlations in Cell Painting compared to L1000, but many compounds are highly correlated in both assays (**Figure 1C**). Interestingly, we observed that certain compounds contained signal in only one assay or the other (**Figure 1C**). We observed that 11% of compounds in the lowest dose (133 / 1,258) and 34% of compounds in the highest dose (422 / 1,258) produced signal in both assays. Combining both assays together, we found that 62% to 85% of compounds (from lowest to highest dose) produced signal higher than random (**Figure 1D**).

### Analyzing the diversity of perturbed cell states manifesting in gene expression and morphology

While percent replicating captures the proportion of compounds that significantly change cell state, it does not quantify the diversity of those cell states when considering the impact of different compounds. Diversity of cell states is critical for many applications, such as MOA matching as described below, because more diversity indicates more potential for interesting biological findings. For example, quantifying cell state diversity is critical when selecting compounds for inclusion in a screening library, as the goal is typically to maximize phenotypic diversity among the compounds; strategies that reduce redundancy allow inclusion of more diverse phenotypes and are therefore more likely to result in drug discovery pipeline “hits” (Wawer et al., 2014).

To qualitatively assess the diversity of profiles produced by each profiling assay, we applied a unified manifold approximation (UMAP) transform (McInnes et al., 2018). We observed that, in both assays, many compounds form distinct islands that consistently group specific MOAs, while a sizable set of compounds are relatively similar to negative controls (**Figure 2A**). MOAs with higher correlations more often form islands in either assay (**Figures S10** and **S11**). The islands separated more with increasing dose, and we identified similarly distributed clusters using t-distributed stochastic neighbor embedding (t-SNE) (van der Maaten, 2008) (**Figure S12)**. Furthermore, a principal components analysis (PCA) grouped together compounds with low replicate reproducibility, representing drug treatments that failed to have a consistent phenotypic impact (**Figure S13**).

**Figure 2.**
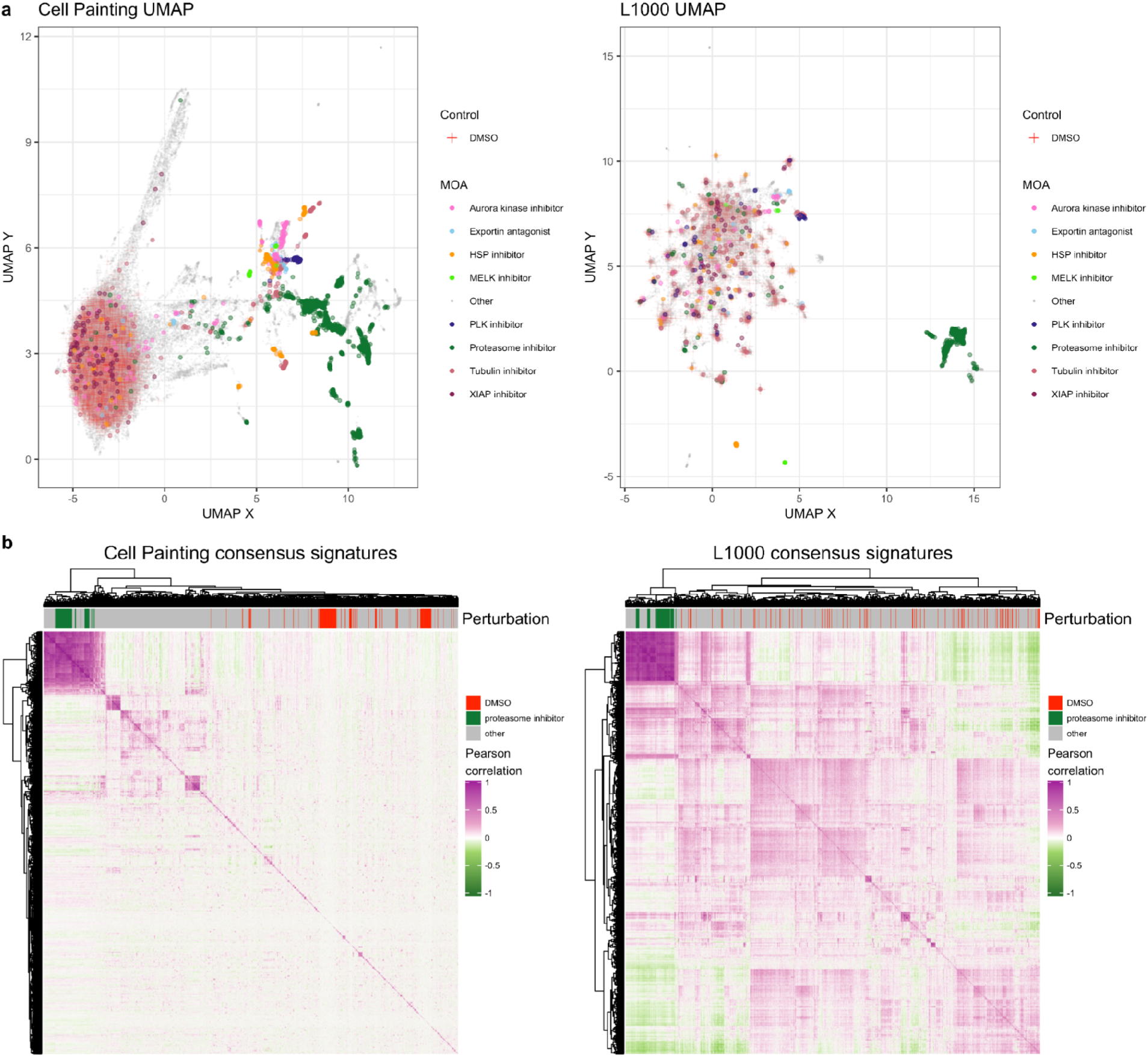
Cell Painting captures a more diverse sample space than L1000. **(a)** Uniform manifold approximation (UMAP) coordinates of all perturbations (level 4 replicates) across all doses by Cell Painting (left) and L1000 profiles (right). We highlight select MOAs that are consistently different from DMSO controls in either modality. Note that Cell Painting data is spherized and L1000 data is not, as explained in the main text; here this manifests in quite different patterns for the negative control DMSO samples. In particular, many of the otherwise-distinct islands of compounds for L1000 are populated by negative control DMSO. **(b)** Heatmaps of pairwise Pearson correlations of all measured compounds’ consensus signatures (see Methods) in either assay at the highest dose (10 μM) plus positive-control proteasome inhibitors at 20 μM and DMSO negative controls.

The primary data collection for this project used a single cell line, A549; a small dataset we gathered using three cell lines (A549, MCF7, and U2OS) showed more phenotypic separation according to cell line and incubation period (**Figure S14**). This separation demonstrated higher biological diversity induced by inherent cell line differences as compared to diversity induced by different perturbations, which is consistent with observations for L1000 data (Squires et al., 2020).

For a quantitative analysis, we fit different clustering algorithms to approximate the number of unique groups of compounds that manifest in each data modality. We observed more distinct clusters in Cell Painting compared to L1000 readouts. This observation was consistent across different clustering solutions (from k=2 to k=40), with different clustering algorithms (k-means clustering and Gaussian mixture models) and using three different metrics (Silhouette scores, Davies Bouldin scores, and Bayesian information criterion) (**Figure S15**). Observing global patterns of pairwise sample correlations in a heatmap provides further evidence of increased diversity in Cell Painting measurements as indicated by lower pairwise correlations across different compounds (**Figure 2B**). Taken together, this analysis suggests that morphology profiles measured by Cell Painting capture more diverse cell states than the gene expression profiles measured by L1000, under the experimental conditions tested.

### Assessing the complementarity of profiling morphology and gene expression features

By design, different profiling technologies measure different biological features. L1000 is a gene expression assay and therefore measures molecular features; specifically, mRNA transcript levels in a biological sample. Cell Painting is an image-based assay that instead measures cell features; both morphological and spatial. Nevertheless, biological signals are often related or even tightly coupled. We therefore sought to approximate how many independent groups of features exist in both modalities. This is distinct from our analysis of the number of groups of *samples* described in the prior section.

In general, we observed higher diversity of feature signals in L1000 compared to Cell Painting (**Figure 3A**). Across doses, individual Cell Painting features had higher coefficient of variance (CV) than L1000 features, but both assays had similar feature variance between replicates (**Figure S16**). Much higher absolute value pairwise correlations among Cell Painting features, even after feature selection (see Methods), indicate more redundant measurements compared to L1000 (**Figure 3B)**. Indeed, the top Principal Components (PCs) explain a higher proportion of variance in Cell Painting compared to L1000 data, providing further evidence of increased feature redundancy in Cell Painting (**Figure 3C**). Both assays attempt to reduce redundancy in some way. For L1000, scientists deliberately chose the genes’ mRNAs to measure (the 978 distinct molecular entities) to minimize redundancy in measurements while maximizing the ability to infer transcriptome-wide gene expression (Subramanian et al., 2017a). Following the standard image-based profiling pipeline (Caicedo et al., 2017), we also removed highly correlated features from the Cell Painting assay. Taken together, this analysis suggests that there is a higher diversity of gene expression features than morphology features, as measured by these two assays.

**Figure 3.**
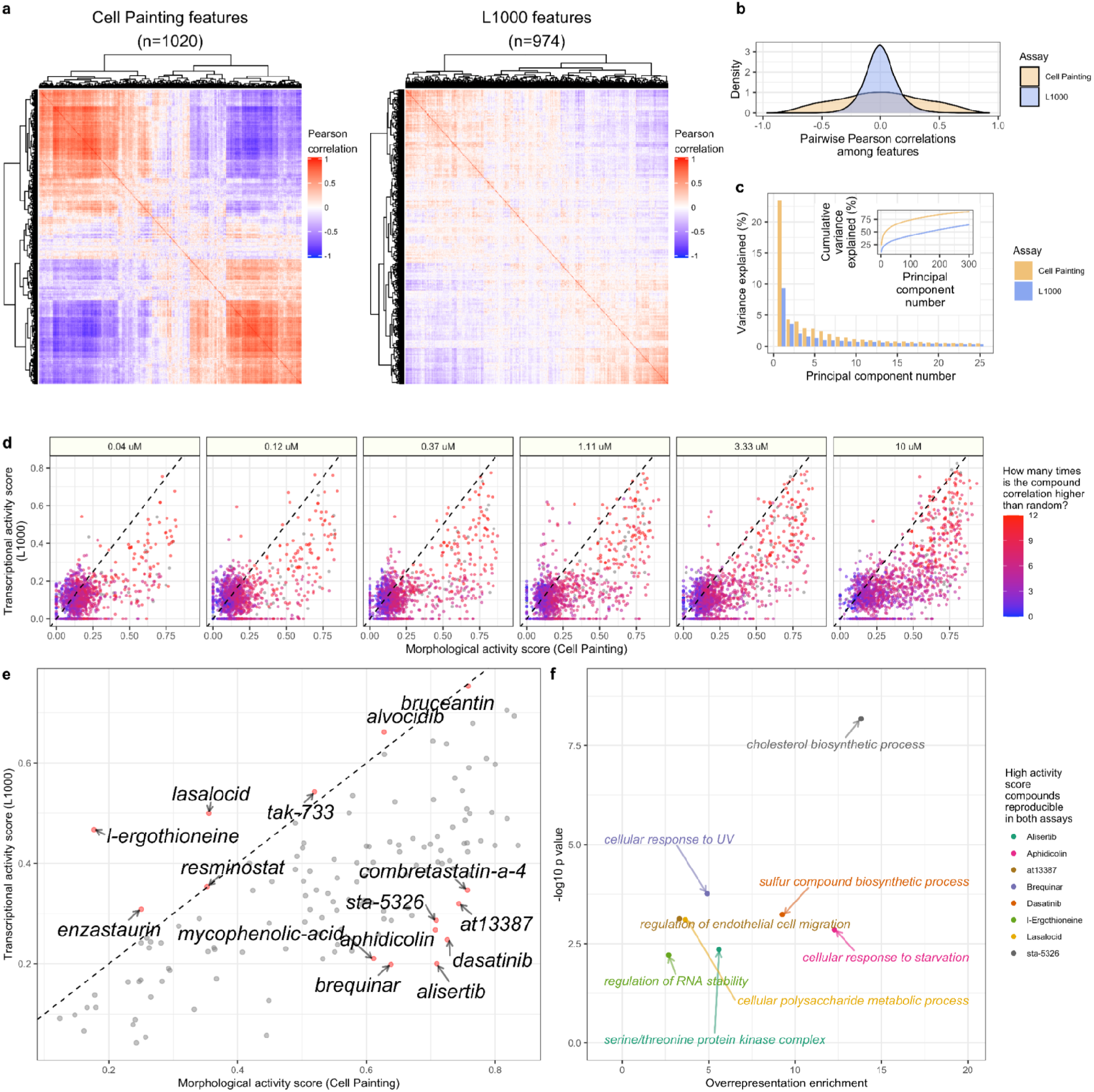
Cell Painting morphology features are more redundant than L1000 gene expression features. **(a)** Heatmaps of pairwise Pearson correlations of 1,020 Cell Painting features and 974 L1000 features, in each case derived from feature-selected consensus signatures of the same compound treatments at 10μM. **(b)** The same data plotted as a density plot shows the distribution of correlations between pairs of L1000 or Cell Painting features. **(c)** The percentage of variance explained for the top 30 principal components derived from a Principal Component Analysis (PCA) in Cell Painting or L1000 readouts. **(d)** Comparing activity scores for highly reproducible compound perturbations (as defined by having 3 or more doses passing the percent strong threshold) reveals that most compounds induce a higher number of morphological changes than gene expression changes. **(e)** The mean MAS and TAS for compounds that are reproducible in at least three doses, with labels for compounds with the largest difference between MAS and TAS. **(f)** Overrepresentation analysis (ORA) for gene ontology (GO) terms using the genes most impacted by each individual compound treatment. We selected these compounds to include those that are reproducible in both L1000 and Cell Painting and that induce a high activity score in one assay, and a low activity score in the other. Each point is a GO term, comprising L1000 landmark genes that were consistently modulated by that compound.

Because we collected thousands of perturbations with replicates, we can study the specific features, in either assay, that were highly impacted by individual compound treatments. Calculating a metric called “activity score” (Subramanian et al., 2017a), which combines both replicate reproducibility and number of impacted features (see Methods), we observed that certain compounds disrupt L1000 and Cell Painting features with different strengths in a dose-dependent manner (**Figure 3D**). Nearly all of the compound perturbations disrupted more morphology readouts than expression readouts. This observation was even more pronounced when we focused on compounds with high reproducibility scores in both assays in at least three different doses (**Figure 3E**). In particular, Dasatinib, Alisertib, Brequinar, Aphidicolin, AT13387, and STA-5326 consistently induced many morphological changes while changing relatively few expression values. By performing separate pathway analyses using the few genes most disrupted by each of the aforementioned compounds, we observed compound-specific associations with specific pathways: Dasatinib altered genes associated with sulfur compound biosynthetic process (GO:0044272, p = 5.8×10^−4^), Alisertib with serine/threonine protein kinase complex (GO:1902554; p = 4.4×10^−3^), Brequinar with cellular response to UV (GO:0034644, p = 1.7×10^−4^), Aphidicolin with cellular response to starvation (GO:0009267, p = 1.4×10^−3^), AT13387 with regulation of endothelial cell migration (GO:0010594, p=7.4×10^−4^), and STA-5326 with cholesterol biosynthetic process (GO:0006695, p =6.7×10^−9^) (**Figure 3F**). Conversely, l-Ergothioneine and Lasalocid induced many more transcriptional changes than morphological changes, which included genes associated with regulation of RNA stability (GO:0043487, p = 6.1×10^−3^) and cellular polysaccharide metabolic process (GO:0044264, p = 7.8×10^−4^), respectively (**Figure 3F**). We provide pathway analysis results for all eight of these high differential activity score compounds in **Table S3**. This type of analysis opens the door to exploring relationships between particular mRNA levels and specific morphologies when perturbing cells (Haghighi et al., 2021; Nassiri and McCall, 2018).

### Assessing the ability of Cell Painting and L1000 to capture compound mechanism of action

We next tested a more demanding, application-oriented metric based on a common use case when profiling compounds: determining a compound’s MOA. A large range of perturbation experiments have mechanistic prediction as a central goal (Schenone et al., 2013). As described in more detail in the Discussion, this is a “notoriously” challenging step in drug discovery where existing methods are useful but have low success rates, usually so low that they have not been calculated systematically. The most common strategy in the pharmaceutical industry is to attempt several painstaking methods and combine results to formulate hypotheses for further testing. In fact, because determining a compound’s mechanism is often time- and labor-intensive, existing annotations for a compound may be incomplete, incorrect, or ignore off-target effects and polypharmacology (Proschak et al., 2019; Rastelli and Pinzi, 2015). Nevertheless, MOA prediction is one of the few biological applications where any modicum of sufficient “ground truth” is available to test a variety of compounds from diverse classes and perform a relative comparison of profiling methods; thus we use it here despite its limitations.

We introduced the metric “percent matching” to quantify how often a profiling assay can group together compound profiles that have the same annotations (see Methods). Unlike percent replicating, this metric is not influenced by plate layout effects in our experiment, because compounds with the same annotated MOA are not located in the same well location across plate maps.

Comparing MOAs within dose, we observed higher percent matching scores for Cell Painting (ranging from 16 - 28%) than for L1000 (7 - 21%) (**Figure 4A**). However, when we compared MOAs across doses, we observed substantially higher percent matching scores for both Cell Painting (44%) and L1000 (50%) (**Figure 4A**). The increased scores highlight the challenges of drug discovery, as many compounds may have different effects at varying doses. Comparing percent matching scores between assays, we observed many overlapping, but also many assay-specific MOAs (**Figure 4B**). In general, we observed stronger signals in L1000 from a smaller number of MOAs, compared to weaker signals from a larger number of MOAs for Cell Painting, as indicated by more points above the dotted line for L1000 but higher percent matching scores for Cell Painting (**Figure 4B** **and** **4C**).

**Figure 4.**
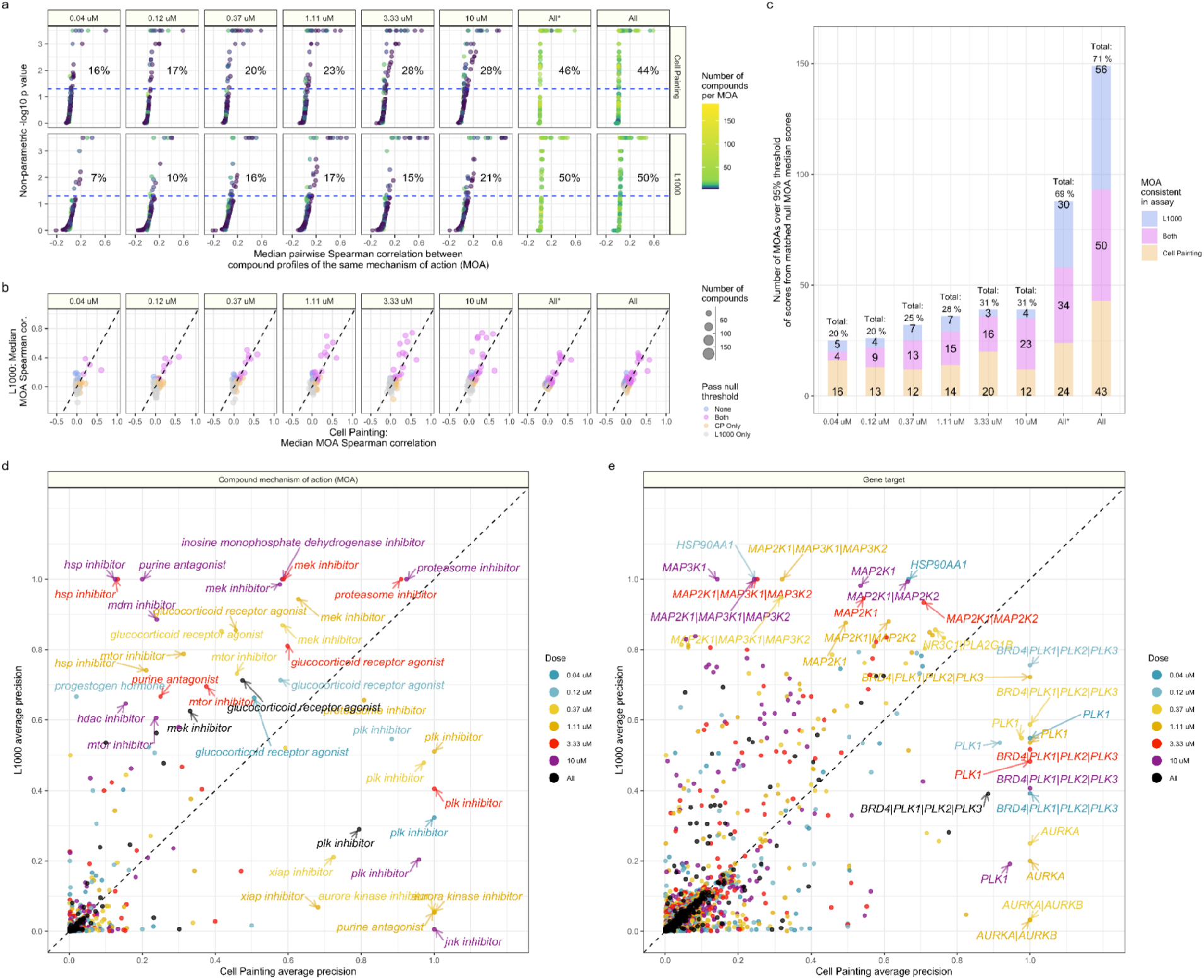
Cell Painting and L1000 differentially measure compound perturbations by mechanism of action (MOA). **(a)** Percent matching metrics for median pairwise replicate correlations of groups of compounds with a given MOA annotation, measured in both assays and across doses. The color of the point represents how many compounds were annotated to a given MOA class. **(b)** Median correlation between compounds annotated with the same MOA. We derived the null threshold through a nonparametric permutation test of randomly sampled compounds (see Methods). The size of the points represent how many compounds belong to the MOA class. **(c)** The L1000 and Cell Painting assays reproducibly measure a complementary set of MOAs. The three numbers represent (from top to bottom) the number of MOAs unique to L1000, the number of MOAs captured in both assays, and the number of MOAs unique to Cell Painting that have higher signal than a randomly permuted null distribution control. The All* bar represents matched MOAs for the 127 MOA set and the All bar represents matched MOAs for the 210 MOA set. Average precision of Cell Painting and L1000 compounds with different **(d)** MOA and **(e)** gene target annotations. We highlight certain high performing MOAs and targets.

The overlapping MOAs, captured by both L1000 and Cell Painting and including at least three different compounds, ranged from 3% of MOAs at the lowest dose (4 / 127), 18% of MOAs at the highest dose (23 / 127), and 27% of MOAs across all doses (34 / 127). Moreover, when considering both assays together, they collectively captured 20% of MOAs at the lowest dose (25 / 127), 31% of MOAs at the highest dose (39 / 127), and remarkably, 69% of MOAs when comparing across doses (88 / 127), although we caution that percent matching scores cannot be directly interpreted as the ability to accurately predict MOA (see next section) (**Figure 4C**). Assaying multiple doses, one captures 19% more MOAs by adding Cell Painting to L1000 and 24% more MOAs by adding L1000 to Cell Painting. These observations can guide researchers in selecting a particular profiling modality that provides more consistent measurements when studying specific compounds or MOAs (**Figure S17A**).

Because percent matching, which is based on the statistical concept of recall, will not sufficiently address how distinguishable MOA classes are from each other (see Methods for more details), we also calculated average precision for all MOAs. Many MOAs demonstrated high average precision across assays and doses, including proteasome inhibitors, MEK inhibitors, and glucocorticoid receptor agonists, but many MOAs were assay-specific (**Figure 4D**). MOA average precision correlated strongly with median pairwise Spearman correlations of MOAs (**Figure S17B**). While we observed increased percent matching when using all doses, we did not observe corresponding increases in all dose average precision, indicating an increase in false positive rate when all doses are considered. Therefore, we advise a careful consideration of both metrics when defining thresholds for follow up experimentation.

We note that in any MOA analysis, low matching scores may result from noise or technical limitations of the assays, but they may also reflect real biological signals resulting from either inaccurate annotations, which is a known challenge (Lin et al., 2019), or alternatively because the assay is capturing mechanistic differences between compounds that are annotated with the same MOA; such polypharmacology is common. We directly observed this difficulty in matching MOAs, as we failed to reliably measure between 102 to 88 different MOAs (80.3% to 69.3% from lowest to highest dose) (**Figure S17C**). Across all comparisons, we failed to reliably measure 29 different MOAs (23%), some of which related to bacterial or fungal processes and others to functions of specialized cell types (**Figure S17D**).

Repeating the average precision analysis using compound gene targets (instead of MOA classes) also revealed high complementarity, as evidenced by many gene targets with off-diagonal precision (**Figure 4E**). L1000 captures activity of compounds targeting MAPK family genes and heat shock protein (HSP90AA1) strongly, while Cell Painting captures aurora kinase genes (AURKA, AURKB), PLK genes (PLK1, PLK2, PLK3) and BRD4 with high precision (**Figure 4E**). We provide a full list of median pairwise replicate correlations, percent matching p values, and average precision metrics for compound MOAs in **Table S4** and compound targets in **Table S5**.

### Different profiling modalities provide complementary deep learning predictions of compound mechanisms and gene target pathways

We next took a targeted approach and trained supervised machine learning algorithms to directly predict compound MOA and gene targets annotated to Gene Ontology (GO) terms. Repurposing the top model architectures from a related Kaggle competition to predict MOA from L1000 readouts and cell viability data (Kaggle.com et al., 2020), we retrained four deep learning models (feed forward neural network, ResNet, TabNet, and a 1D convolutional neural network) and a K-Nearest Neighbor baseline on multi-label objectives. We also averaged model probabilities to form an ensemble model, and considered compounds that map to multiple MOAs and gene targets as positive labels for all individual categories. We compared model performance for training each model using L1000 and Cell Painting readouts separately and merged together (**Figure 5A**).

**Figure 5.**
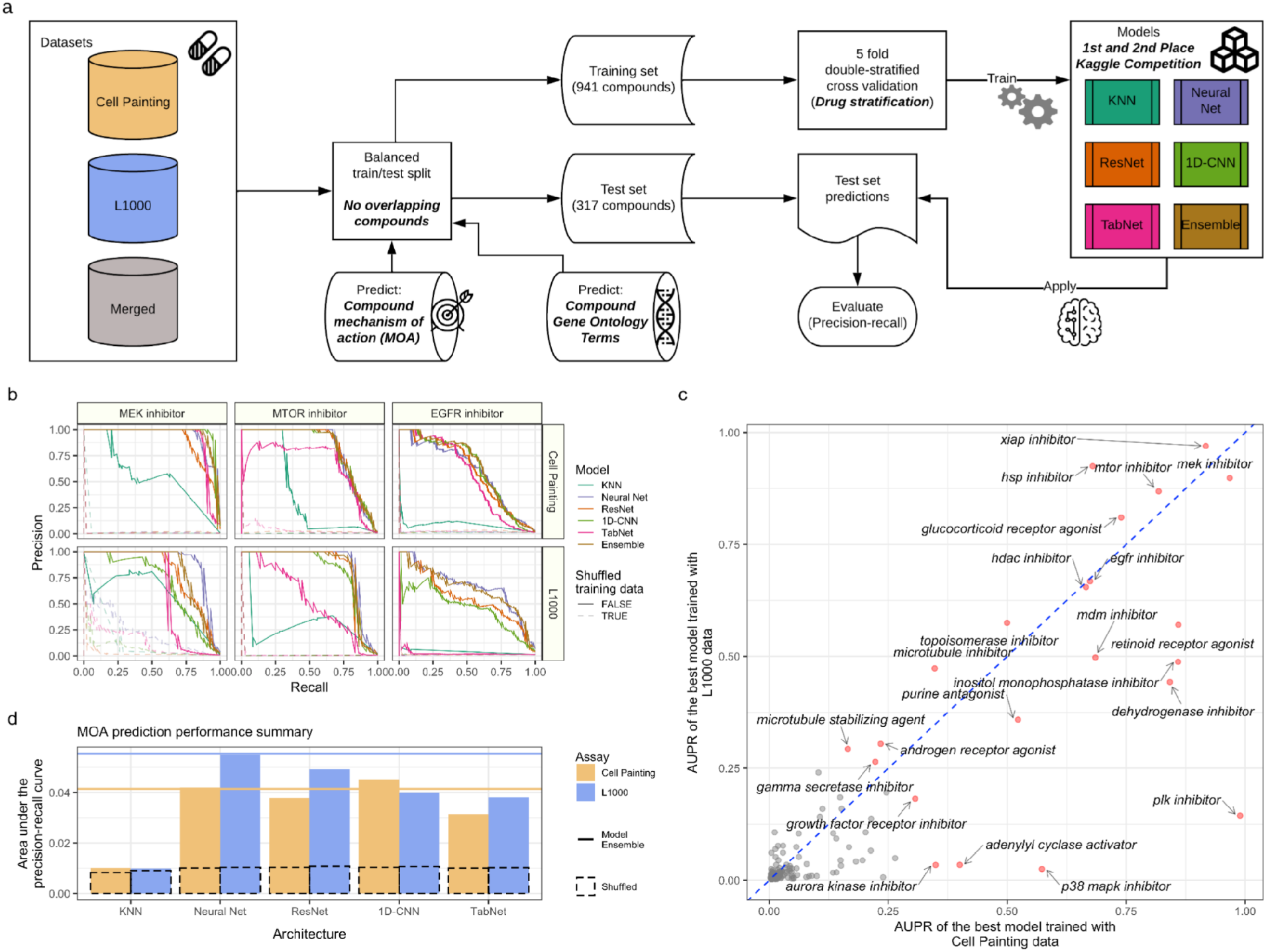
Predicting compound mechanisms of action (MOA) in Cell Painting and L1000 reveals overlapping and complementary performance for different mechanisms. **(a)** Deep learning workflow. We collected compound Cell Painting and L1000 data from compound perturbations and trained five different deep learning models to predict compound MOA and Gene Ontology terms. **(b)** Held out test set precision-recall curves for three well performing MOAs in both assays. **(c)** Individual MOA performance by held out test set area under the precision-recall curve (AUPR) in the top performing model using Cell Painting and L1000 data. **(d)** Overall held out test set model performance measured by AUPR for MOA prediction for our multi-label, multi-class prediction framework. We trained models from a recent Kaggle competition plus a K nearest neighbors baseline model. The dotted bar chart represents a negative control in which we trained models with shuffled labels. The solid lines indicate ensemble model performance by blending model predictions (see Methods). We trained all models using level 4 replicate profiles.

Many individual MOAs could be predicted rather robustly by both assays including MEK inhibitors, MTOR inhibitors, and EGFR inhibitors (**Figure 5B**). However, some MOAs could be consistently predicted better by L1000 (e.g. HSP inhibitors) or by Cell Painting (e.g. PLK inhibitors), and many MOAs could be predicted by neither assay (e.g. Glucocorticoid receptor antagonist) (**Figure S18A**). In the held out test set, the MOA predictions were correlated between the two assay modalities (Spearman correlation = 0.70, p < 2.2e-16), but some MOAs were predicted better in one assay compared to the other (**Figure 5C**). In general, while performance across all MOA predictions sufficiently improved over several baselines, overall performance was relatively low, demonstrating the general difficulty of the task (**Figure 5D**). Poor predictions might be a result of noisy readouts or the ability of the data type to reveal more subtle compound-specific signals such as off-target effects.

Overall, L1000 performed slightly better at MOA prediction than Cell Painting across a wide range of different deep learning architectures (**Figure 5D**). The Kaggle competition selected for models especially suited to L1000 and cell viability data; it is possible that alternate architectures might favor Cell Painting data. Concatenating features of both assays together and predicting MOA did not improve performance, but this is likely because of the need to randomly match replicates since there is no one-to-one correspondence (see Methods) (**Figure S18B**). Subsampling the Cell Painting data to match the sample count of L1000 slightly decreased performance, indicating that collecting more than three replicates in either assay is likely to increase performance (**Figure S18B**). We also observed that performance increased in step with treatment dose (**Figure S18C**).

In addition to MOA predictions, we also trained these same deep learning architectures to predict gene pathways. The genes that compounds target are often well characterized, and we hypothesized that Cell Painting and L1000 could predict compounds targeting genes in the same pathway. Therefore, we mapped Drug Repurposing Hub compound gene target annotations to Gene Ontology (GO) terms, and retrained the same MOA models to instead predict GO terms (Ashburner et al., 2000; Gene Ontology Consortium, 2021). In our dataset, we assayed compounds that targeted 720 different genes, which mapped to 5,822 unique GO terms. After filtering GO terms that included 20 different compounds or more, we trained models to predict different terms.

With slightly worse performance than the MOA predictions, many individual GO terms could be predicted by either assay including neuron differentiation (GO:0030182), chromatin organization (GO:0006325), and steroid hormone binding (GO:1990239) (**Figure S19A**). As expected, some GO terms were better-predicted using L1000 data (e.g. transmembrane receptor protein tyrosine kinase activity; GO:0004714), others with Cell Painting data (e.g. positive regulation of G1/S transition of mitotic cell cycle; GO:1900087), and others by neither assay (e.g. Cellular response to calcium ion; GO:0071277) (**Figure S19B**). However, GO term prediction performance was highly correlated between the two assays (Spearman correlation = 0.92, p < 2.2e-16; **Figure S19C**). We observed improvements over baselines, but general poor performance overall (**Figure S19D**). We provide all deep learning performance metrics in **Table S6**.

## Discussion

Large-scale perturbational profiling experiments are time- and cost-intensive; comparing their relative abilities is important information for experimental design and planning. We found that mRNA profiling (via L1000) and morphological profiling (via Cell Painting) were generally complementary, given their current state of technical reproducibility and standard computational pipelines. Cell Painting had a higher diversity of samples and could match MOAs more consistently in an unsupervised setting, while L1000 had a higher diversity of features and better performance predicting MOAs in a supervised setting. Cell Painting is less expensive, enabling larger experiments for a given budget, but L1000 offers a larger pool of publicly available data to query (Subramanian et al., 2017a). A wide variety of biological pathways are readily captured by both data types, but some are better observed in one modality versus the other.

We had anticipated that morphological changes would generally not occur without concomitant changes in mRNA levels (whether as a cause or consequence), particularly at low doses, but found examples of compound treatments where this happens, and vice versa. It may be that the L1000 assay does not capture the mRNAs in the cell that change, due to technical noise or because it measures only ~5% of the transcripts in the cell. It could also be that mRNA changes do occur, but not at the short timescale of the mRNA detection (24 hours) as compared to the image capture (48 hours). It is also possible that the different incubation times for compound treatments increased our ability to detect changes, but it is unlikely any single time point is optimal for all treatments, given potential differences in cell and molecular response times (Niepel et al., 2017).

MOA prediction is “notoriously challenging” and a major bottleneck (Lill et al., 2021). No assay exists that can reliably succeed across a majority of known MOA classes (Pasquer et al., 2020). Of course, one would not expect any assay in a single cell line at a single time point or compound dose, to capture 100% of all biological mechanisms. For this reason, in practice it is more typical to assess compounds of interest with various strategies, each with low individual success rates. Our goal in this analysis was not to prove the efficacy of MOA prediction by mRNA and morphology profiling. Instead, in the MOA prediction analyses, we aimed to compare the relative strengths of the two assays, because this is a direct comparison task with the most available ground “truth”. In fact, both assays tested here have the advantage of being sufficiently inexpensive; most other strategies for MOA prediction report no success rates because they are not practically scaled to test on thousands of compounds systematically, for example, those that require modification of the compound or customization of cells (Pasquer et al., 2020).

One can imagine future developments in both assays that could improve their performance in a variety of applications. For perturbational profiling of mRNA expression, the L1000 assay offers a cost effective strategy for large-scale compound experiments, with a huge library of publicly available profiles. Fortunately, expression profiles from L1000 are similar to RNA-seq equivalents: out of 3,176 patient-derived RNA samples profiled on both platforms, 3,103 (98%) had high quality cross-platform correlations (Subramanian et al., 2017b). The development of novel methods that are even cheaper, more robust, and more comprehensive would be welcome. For imaging assays, deep learning-based segmentation and feature extraction offers promise but deep learning is not yet routine for image-based profiling (Chandrasekaran et al., 2021; Pratapa et al., 2021). As well, the standard Cell Painting pipeline population-averages measurements; methods that better leverage the assay’s single-cell measurements are likely to improve information capture from this assay. Using additional stains is another sensible route, although initial testing indicates it does not seem to dramatically improve MOA prediction performance (Rose et al., 2018). For both, screening additional cell types (Boyd et al., 2019; Cox et al., 2020; Rose et al., 2018) and timepoints might increase the ability to detect and characterize perturbations in cell state. If experiments capture both profiling types, the profiles might be integrated to increase their power and resolution (Caicedo et al., 2021b; Haghighi et al., 2021; Huang et al., 2017; Lapins and Spjuth, 2019). Overall, this paper will help researchers to better understand the pros and cons of the two currently largest and cheapest methods of large-scale drug profiling.

## Acknowledgements

The authors thank Joshua Sacher for his help in curating Drug Repurposing Hub compound metadata. The authors gratefully acknowledge funding from the National Institutes of Health (NIH R35 GM122547 to AEC), including the Common Fund LINCS Grant (NIH U54 HG008699 to AS) as well as an internship funded by the Massachusetts Life Sciences Center (to AA). We would also like to acknowledge the use of the Opera Phenix High-Content/High-Throughput imaging system at the Broad Institute, funded by the S10 Grant NIH OD-026839-01.

## Author contributions

G.P.W., T.N., S.S., A.S., and A.E.C. conceived the study. G.P.W. and T.N. performed the bioinformatics and data processing pipelines. G.P.W., T.N., A.A., L.L., A.X.Y., J.C.C., M.H.R., M.H., and S.S. performed analytical experiments. B.A.C., K.K., and D.J.L. designed and performed the image analysis pipelines. M.K.A and K.H. performed the microscopy experiments. X.L. performed the L1000 experiments. G.P.W., A.A., M.B., B.A.C., S.N.C., E.W., and S.S. made key software contributions to curate and process data. G.P.W., T.N., S.S., A.S., and A.E.C. drafted and edited the manuscript. All authors approved the final version.

## STAR Methods

### LEAD CONTACT AND MATERIALS AVAILABILITY

***Lead Contact:*** Further information and requests for resources should be directed to and will be fulfilled by the Lead Contact, Anne Carpenter.

#### Data and code availability

##### Source data statement

All code to reproduce this analysis is located at https://github.com/broadinstitute/lincs-profiling-complementarity. All code to reproduce the Cell Painting image-based profiling pipeline is available at https://github.com/broadinstitute/lincs-cell-painting. The L1000 data are available at figshare. Cell Painting images are deposited to the Image Data Resource (https://idr.openmicroscopy.org/) under accession number idr0125. Cell Painting images and single-cell profiles are available at the Cell Painting Gallery on the Registry of Open Data on AWS (https://registry.opendata.aws/cellpainting-gallery/) under accession number cpg0004. This paper also analyzes existing, publicly available data. These accession numbers for the datasets are listed in the Key Resources Table.

##### Code statement

All code to reproduce this analysis is located at https://github.com/broadinstitute/lincs-profiling-complementarity, which we archived on Zenodo (Way et al., 2021a). All code to reproduce the Cell Painting image-based profiling pipeline is available at https://github.com/broadinstitute/lincs-cell-painting, which we archived on Zenodo (Natoli et al., 2021b). For all analyses, we used Python version 3.9.1 (Van Rossum and Drake, 2009) and pandas version 1.2 (McKinney, 2010). For visualization we used R version 3.5.1 (R Core Team, 2021) and ggplot2 version 3.3.0 (Wickham, 2016). For versions of other critical software see our Key Resources Table and as discussed above. All computational environments can be reproduced in our github repository https://github.com/broadinstitute/lincs-profiling-complementarity. We used conda version 4.10.3 and conda-forge to version all computational environments (Anaconda Inc., 2021; Community, 2015)

Any additional information required to reanalyze the data reported in this paper is available from the lead contact upon request.

### EXPERIMENTAL MODEL AND SUBJECT DETAILS

#### Selection of compounds for testing

We selected compounds annotated by the Drug Repurposing hub using two criteria: 1) Compounds known to have diverse phenotypic outcomes with a variety of annotated MOAs; 2) Compounds that are in current clinical use or in advanced clinical testing (we deprioritized tool compounds). We did not filter our compound selection for prior performance-based observations based on past datasets.

We exposed A549 cells to six different doses of most compounds. The six dose points include 0.04μM, 0.12μM, 0.37μM, 1.11μM, 3.33μM, and 10μM. For several compounds the calculated dose was slightly different than one of the six dose categories, and in these instances, we rounded to the nearest dose category. The complete list of all compounds tested and their annotations can be found in **Table S1**.

### METHOD DETAILS

#### Sample preparation: Cell Painting

We generated Cell Painting data according to (Bray et al., 2016). Briefly, we cultured A549 cells in RPMI (Mediatech) on 384 well plates and exposed them to compound treatment at various doses for 48 hours. After exposure, we fixed, stained, and then imaged all cells. Specifically, we used Hoechst 33342 to mark DNA, concanavalin A/Alexa Fluor 488 conjugate to mark the endoplasmic reticulum (ER), SYTO 14 to mark nucleoli and cytoplasmic RNA, phalloidin to mark F-actin cytoskeleton, wheat-germ agglutinin/Alexa Fluor 555 conjugate (WGA) to mark Golgi and plasma membrane, and MitoTracker Deep Red to mark mitochondria. For complete details about the Cell Painting procedure, see (Bray et al., 2016).

We performed all imaging using a Phenix Opera with a 20X/1.0NA water objective, 1×1 binning, and filter sets described in Bray et al 2016 Supplementary Note 1. For the second batch of Cell Painting data (**Figure S14**) we treated cells at the same doses for 6, 24, and 48 hours.

#### Sample preparation: L1000

We generated the L1000 data according to the protocol outlined in (Subramanian et al., 2017a). Briefly, we cultured A549 cells in RPMI (Mediatech) on 384 well plates and exposed them to compound treatment at various doses for 24 hours. After the incubation time, we lysed cells and subjected them to ligation-mediated amplification (LMA) and detection. We captured mRNA using oligo-dT coated beads and reverse transcribed the sequences into cDNA. We PCR amplified the cDNA using biotinylated, barcoded primers and gene-specific juxtaposed probe pairs resulting in gene-specific, barcoded, and biotinylated PCR amplicons. We then hybridized these amplicons to Luminex beads, stained them with streptavidin R-phycoerythrin (SAPE), and detected them using a Luminex FlexMAP 3D scanner. Therefore, each bead reports the barcode, which determines gene identity, and we measure the SAPE fluorescent intensity, which indicates transcript abundance.

#### L1000 data processing

We processed L1000 data into perturbagen-specific differential expression signatures as described in (Subramanian et al., 2017a). Briefly, we captured raw fluorescent intensities (FI) from the Luminex FlexMAP 3D scanner for each of the 978 L1000 landmark genes (Level 1 data). We then deconvoluted FI data to extract the median FI (MFI) for the two genes being measured by each Luminex bead barcode (Level 2 data). We loess-normalized the MFI values to the ten L1000 invariant gene sets within each well, and then quantile normalized all wells on the same detection plate, which resulted in each sample having the same empirical distribution (Level 3 data). We then computed gene-wise robust z-scores per sample, using all other samples on the same plate as the reference distribution (Level 4 data). Lastly, we collapsed biological replicates into consensus signatures using a weighted average, where each replicate was weighted by its average correlation with the others (Level 5 data). We made all data and metadata publicly available on figshare (Natoli et al., 2021a).

#### Image feature extraction

To extract image features, we built a CellProfiler (version 2.3.1) (Kamentsky et al., 2011) image analysis pipeline and ran it on Amazon Web Services using Distributed-CellProfiler (McQuin et al., 2018). We also performed illumination correction to standardize readouts and account for confounding factors by homogenizing light across all fields of view (Singh et al., 2014). The image analysis pipeline segments cells by distinguishing nuclei from cytoplasm and then extracts measurements for specific features related to the various channels captured (see Sample preparation: Cell Painting). Specifically, we measured fluorescence intensity, texture, granularity, density, location, and various other measurements for each single cell (see https://cellprofiler-manual.s3.amazonaws.com/CPmanual/index.html for more details). Following the image-analysis pipeline, we obtain 110,012,425 cells and 1,790 feature measurements across 136 different plates. The pipelines are available online here https://github.com/broadinstitute/imaging-platform-pipelines/tree/3eb4ff5676aa7889666f09b606cd915c8b9ea839/cellpainting_a549_20x_phenix_bin1.

#### Cell Painting image-based profiling

After extracting CellProfiler readouts from all Cell Painting images of segmented single cells, we applied an image-based profiling pipeline to process morphology readouts (**Figure S3B**). In the first step of this pipeline, we used cytominer-database (https://github.com/cytomining/cytominer-database) to collect and validate all CellProfiler output measurements from Cells, Cytoplasm, and Nuclei compartments for every site (field of view). The output of this first step is a set of SQLite files that contain raw single cell profiles per plate (level 2 data).

Next, we used pycytominer to process the single cell readouts (Way, G.P., Chandrasekaran, S.N., Bornholdt, M., Fleming, S.J., Tsang, H., Adeboye, A., Cimini, B., Weisbart, E., Ryder, P., Stirling, D., Jamali, N., Carpenter, A.E., Singh, S., 2021). We performed a standard image-based profiling pipeline (Caicedo et al., 2017) consisting of profile aggregation, annotation (level 3 profiles), normalization (level 4a), feature selection (level 4b), and forming consensus signatures (level 5). We performed median aggregation and normalized aggregated profiles using the “mad_robustize” method, which scales features independently by subtracting each value by the median and dividing by the median absolute deviation. We normalized each plate using the DMSO controls only, which allows us to more easily compare profiles across plates. We also performed several standard feature selection operations to remove features with missing data (“drop_na_columns”), remove features with low variance (“variance_threshold”), remove features that are highly correlated with other features (“correlation_threshold”), and remove blocklist features (“blocklist”). These blocklist features include CellProfiler features that we’ve previously observed to be unstable and noisy (Way, 2020).

Because the negative control DMSO profiles were noisy due to technical artifacts, we applied a spherize transform (also known as whitening) to mitigate the impact of well positioning (Ando et al., 2017; Kessy et al., 2018). More specifically, we used the zero-phase whitening filters (ZCA) solution calculated on the profile correlation matrix (ZCA-cor) to minimize the absolute distance between the transformed profiles and the untransformed profiles (Bell and Sejnowski, 1997). We also formed consensus signatures (level 5) by moderated z-score (MODZ) aggregating all replicate wells across plate maps into a single signature. We applied feature selection to the consensus signatures and batch effect corrected profiles separately using the same operations as described above. We applied the same pipeline to batch 1 (A549) and batch 2 (A549, MCF7, and U2OS) Cell Painting datasets.

Different drug treatments induce differing amounts of cell death and cell growth rates. To predict the amount of cell death, we applied a recently derived machine learning model to predict cell death readouts from Cell Painting features (Way et al., 2021b) We specifically used the “percent dead” machine learning model originally trained using the cell viability panel of the Cell Health assay to make predictions (see **Figure S4**).

We provide all the image-based profiles (level 3 and up) and the data processing pipelines in a versioned and publicly available github repository at https://github.com/broadinstitute/lincs-cell-painting/ (Natoli et al., 2021b).

#### Calculating reproducibility metrics - percent replicating

The first step in calculating percent replicating is to calculate the median pairwise Spearman correlation of all treatments (compound and dose). We determined if this median correlation was greater than what we expected by chance by comparing it to carefully-matched null distributions. We report the metric per assay (under different normalization and null distribution conditions). See **Figure S6** for a graphic fully explaining this metric, and see below for a full description of how we designed the null distribution.

We designed null distributions to control for three things: 1) different replicate cardinalities between different compound treatments, 2) well position on the 384 well plate, and 3) treatment dose. We controlled for replicate cardinalities to account for stability in median values across sample counts, the position of the well on a plate to account for potential plate position effects, and treatment dose to account for the higher likelihood that higher doses contain more non-specific signals, and would therefore result in higher absolute correlations between unrelated compounds.

Specifically, for percent replicating, for a given perturbation *x* located on well *w* measured across *n* replicates and treated with dose *p*, we randomly sampled *n* non-replicate profiles assayed in well *w* (but from different plate maps) from all perturbations that were treated with dose *p*. We performed this sampling procedure 1,000 times per replicate cardinality (e.g. compounds with 3 replicates, 4 replicates, 5 replicates, etc.) with two additional restrictions: (1) the random sample did not include replicates for perturbation *x*, and (2) no two compounds of the same non-*x* perturbation were included in the same null group. For example, in cases where a compound treatment at a specific dose had five replicates, we sampled 1,000 groups of five randomly sampled non-replicate profiles of the same dose. We used level 4 profiles considering compound and dose information as replicates, and we considered a replicating profile one in which the ground truth median pairwise replicate correlation was higher than 95% of the null distribution. We therefore calculate the percent replicating metric as the proportion of all replicating profiles over all common perturbations. This 95% thresholding procedure is equivalent to calculating per-treatment non-parametric p values (by counting how many times the replicate pairwise correlation was greater than the non-replicate null distributions) and reporting how many compounds were above an alpha p-value threshold of 0.05. We report this percent replicating implementation in **Figure 1**.

We also calculated percent replicating by relaxing the two null distribution constraints separately. We performed the procedure as described above except we 1) did not require the non-replicates be drawn from the same well position and 2) did not require the non-replicates to be drawn from the same dose. We relaxed these constraints to observe the impact of well position and dose on percent replicating interpretation. We compare these results in **Figure S7**.

#### Calculating reproducibility metrics - percent strong

We also introduced and calculated a second metric, which we called “percent strong” (see **Figure S8**). In percent strong, we construct the non-replicate null distribution without adjusting for well position or replicate cardinality. We did still, however, calculate both dose-specific and dose-independent null distributions. Specifically, for dose-specific metrics, for each modality and normalization strategy independently, we calculate a single null distribution for each dose by randomly sampling 1,000 groups of non-replicate profiles per replicate cardinality (the same null distributions for percent replicating, but we ignore replicate cardinality) and compare them to median non-replicate pairwise correlations. For our dose-independent analysis, we do not restrict profiles from being measured at the same dose.

We subsequently calculate percent strong as the percentage of replicate median pairwise Spearman correlations greater than 95% of the full non-replicate null distribution. Percent strong provides more possible combinations of non-replicate sampling and therefore is not as susceptible to sampling biases as percent replicating. In other words, because percent replicating strictly samples non-replicates from the same well, if a specific well, by chance, housed similar perturbations, the non-replicate distribution might be unduly skewed and deflate percent replicating scores. Percent strong is the least constrained null distribution and is robust to normalization strategy and subsampling (see subsampling subsection).

We calculated percent replicating and percent strong using Cell Painting and L1000 input data with five different normalization strategies: 1) Cell Painting level 4 spherized profiles; 2) Cell Painting level 4 non-spherized profiles (median aggregated features with z-score normalization); 3) Cell Painting level 4 spherized subsampled profiles (see below); 4) L1000 level 4 spherized profiles; and 5) L1000 level 4 non-spherized profiles. We also calculated percent strong without dose-specific null distributions (dose-ind.) and after filtering edge wells using spherized Cell Painting level 4 and non-spherized L1000 level 4 profiles.

#### Subsampling Cell Painting level 4 profiles to match L1000 replicate count

We collected fewer L1000 profiles than Cell Painting profiles. In most cases, with some exceptions, we collected three L1000 replicates and five Cell Painting replicates. We collected samples according to standard operating procedures for both assays, which pertain to sample handling and costs.

To determine the extent to which our percent replicating metrics were biased by replicate count, we performed a subsampling experiment using the spherized Cell Painting profiles. Specifically, we randomly sampled Cell Painting profiles without replacement to match exactly the same number of L1000 replicates for the individual compound of interest. Using this subsampled dataset, we calculated percent replicating. We also recalculated the null distribution using subsampled profiles.

#### Plate diffusion analysis to test the impact of plate position effects

We performed a plate diffusion analysis to assess plate position biases in Cell Painting and L1000 data. Specifically, for a given well *w* with treatment *x* collected on plate map *P*, we collected all non-replicate samples across all plate maps except *P* in a specific well neighborhood as defined by diffusion parameter *d*. In other words, we selected all non-replicate wells in a predefined local neighborhood around well *w*. We used five different diffusion parameters (0, 1, 2, 3, and 4) to define this neighborhood. For d=0, we only included non-replicate samples from the same well, for d=1, we included all adjacent neighbors of well w on different plate maps, for d=2, we included all adjacent neighbors plus all neighbors’ neighbors on different plate maps, and so on. After defining these non-replicate samples, we calculated all combinations of pairwise replicate correlations between treatment *x* and all non-replicate samples and calculated the mean of the distribution of well-neighborhood pairwise correlations.

Furthermore, we not only considered the local neighborhood around well *w*, but also the local neighborhood around the hypothetical plate-flipped version of well *w* (e.g. well P24 is the flipped version of well A01) in collecting non-replicates to analyze. In practice, the scientists collecting the data put the 384-well plate in the data collection machine in one of two orientations. Including this mirror parameter ensures that our diffusion analysis captures any technical plate effects introduced by different plate orientations.

We used the same five level 4 input data sets with different normalization strategies as we defined in the percent replicating and percent strong methods subsection. We report the mean of the total well-neighborhood pairwise correlations to determine consistent plate position technical artifacts per well position. If a strong plate position effect were present, then we would expect to see neighborhood correlations substantially drop with increasing diffusion.

#### Quantitative assessment of profile clustering

Using spherized Cell Painting level 4 profiles and non-spherized L1000 level 4 profiles, we performed three iterative clustering analyses in which we fit algorithms across a range of cluster numbers between k = 2 and k = 40 and acquired three goodness-of-fit heuristics (Silhouette scores, Davies Bouldin scores, and Bayesian Information Criterion (BIC) scores) for both datasets. Briefly, the Silhouette score is a metric indicating how separable clustering solutions are, with a score of 1 indicating that the identified clusters are clearly separable (Rousseeuw, 1987). The Davies Bouldin score quantifies the ratio of within-cluster distances to between-cluster distances when comparing each cluster to their most similar neighboring cluster, and a lower value indicates more separable clusters (Davies and Bouldin, 1979). BIC is a measurement of cluster likelihood and cluster predictability with an added penalty for increased cluster number, and a lower value indicates better clustering (Schwarz, 1978). We visualize the tradeoff of these heuristics as we fit clustering algorithms with increasing cluster numbers.

For each model fitting, we used all 1,327 common compounds transformed into PCA space using 350 components. Therefore, we fit all clustering algorithms and calculated goodness-of-fit metrics using data of the same feature dimension, which, if not identical, can skew metrics and make comparison difficult.

Specifically, we applied k-means clustering with a maximum of 1,000 random iterations, across the k=2 to k=40 cluster number range, and calculated Silhouette and Davies Bouldin scores from the resulting cluster solutions. We also fit full covariance Gaussian Mixture Models (GMM) with a k-means initialization and 1,000 maximum iterations, and we calculated BIC scores from the resulting clustering solutions. We performed this procedure using profiles resulting from each of the six different treatment doses independently, as well as using all profiles combined.

#### Calculating signature strength and activity score

To compare how different compound perturbations impacted individual feature measurements for both L1000 gene expression and Cell Painting morphology assays, we calculated signature strengths and activity scores as previously described (Subramanian et al., 2017a). Specifically, signature strength counts the number of features that substantially change when a sample is perturbed with a specific compound. We determined a substantially changed feature as one with a value greater than 2 after multiplying its z-score (transformed with respect to all compounds) by the square root of the number of replicates. We multiply by the square root of the number of replicates to enable more direct comparison of scores across compounds with different replicate counts.

Counting features in this fashion is equivalent to computing the absolute magnitude of change – we are implicitly transforming each feature so that values above 2 (or below −2) are mapped to 1 (or −1) and the rest are mapped to 0 (a “hard” sigmoid), and are then measuring the L1 norm (or L1 magnitude) of the resulting transformed vector. Intuitively, compounds that induce many features to high absolute value z scores are disruptive of steady state, and compounds that don’t change many features are not broadly strong perturbations. Instead, these compounds may either have little impact or be highly specific, meaning they only target one, or a few features strongly.

Activity score, either Morphological Activity Score (MAS) or Transcriptional Activity Score (TAS) for the Cell Painting or L1000 assays respectively, is the geometric mean of signature strength and median replicate correlation, normalized by the square root of number of features in the assay such that the resulting metric ranges between 0 and 1. A high activity score indicates compounds that reproducibly induce large changes in many features for a particular assay readout.

#### Identifying independent groups of features in assay measurements

To analyze feature redundancy and estimate the number of feature modules per assay, we calculated pairwise Pearson correlations of level 5 consensus profiles of Cell Painting (spherized) and L1000 (non-spherized) assays. We applied the same feature selection procedure in both assays, using pycytominer (Way, G.P., Chandrasekaran, S.N., Bornholdt, M., Fleming, S.J., Tsang, H., Adeboye, A., Cimini, B., Weisbart, E., Ryder, P., Stirling, D., Jamali, N., Carpenter, A.E., Singh, S., 2021). Specifically, we removed redundant features (as defined as having pairwise Pearson correlations < 0.9), features with low variance, and blocklist features (Way, 2020). This resulted in 1,020 Cell Painting features and 974 L1000 features. We calculated pairwise Pearson correlations of these features for all common compounds perturbed with 10 μM of compound. We visualized feature-level correlations using ComplexHeatmap (Gu et al., 2016).

Using sci-kit learn (Pedregosa et al., 2011), we applied principal component analysis (PCA) with n_components = 150 using feature-selected level 4 profiles for each assay independently. PCA provides the percentage of variance explained for each orthogonal component, and we use this information to determine the variety of signals in each feature space (Jolliffe, 1986).

#### MOA prediction - Calculating percent matching

For our percent matching metric, we performed a similar procedure as percent replicating (see above). The only differences were that we (1) used level 5 consensus signatures from both data sets and (2) considered MOA and dose information as replicates. We used level 5 consensus signatures instead of level 4 replicate signatures, because consensus signatures are less noisy and correct for potential different replicate cardinalities per compound within an MOA. We only considered MOAs that had three or more annotated compounds. This resulted in an analytical set of 127 unique MOAs. We considered compounds annotated with multiple MOAs as independent entities.

We subsequently constructed dose and MOA compound cardinality-specific null distributions to compare against. Specifically, for each MOA, we calculated its median pairwise replicate correlation. We next randomly sampled 1,000 groups of level 5 consensus profiles of the same cardinality of the MOA compound count. For example, if an MOA contained 10 compounds, we formed one group by randomly sampling 10 compounds from different MOAs. We only sampled compounds measured at the same dose, and we did not include any two compounds of the same MOA in each random sample. For each of the 1,000 randomly groups, we calculated median pairwise correlations, which formed our percent matching null distribution. Lastly, we calculated a compound specific p value by dividing how many times the real median pairwise correlation of replicates was higher than all 1,000 randomly sampled groups of median pairwise correlations. We considered a matched MOA one in which the ground truth MOA median pairwise correlation was higher than 95% of the null distribution. We therefore calculate the percent matching metric as the total number of matched MOAs over all common MOAs.

We also repeated the above percent matching procedure without the same-dose requirement for 1) the replicate compounds belonging to the same MOA and 2) the randomly sampled null distribution. To prevent overinflated metrics and to ensure signal is driven by different compounds of the same MOA, we did not consider replicates of the same compound across different doses when calculating median pairwise replicate correlations. In addition to calculating all-dose percent matching with the core 127 MOAs that passed the within-dose compound count filtering criteria, we also calculated percent matching with a relaxed MOA filtering. Specifically, when we considered the “All” dose comparison, we relaxed the MOA count constraint to contain two or more annotated compounds, which resulted in 210 unique MOAs. In effect, this enabled us to compare MOAs with two annotated compounds across multiple doses (“All” vs. “All*” bars in **Figure 4** and **Figure S17**).

#### Calculating average precision for compound MOAs and compound gene targets

In a similar approach as described in the percent matching section above, we also calculated average precision as a metric to compare similarity of profiles targeting the same MOA and the same genes. We used this additional metric because percent matching measures cluster compactness and recall, and thus risks overlooking differences in separation among clusters of samples. Although the degree of separation among clusters is captured to some degree by the null, especially when it includes other samples and not just the negative control DMSO, this does not fully mitigate the issue. By contrast, average precision instead estimates the area under the precision recall curve, by averaging precision at various recall thresholds, thus measuring cluster separation, or, how compounds from the same MOA appear different from other compounds.

We used cmapPy version 4.0.1 to calculate pairwise Pearson correlations of all profiles within each dose and assay, independently (Enache et al., 2018). To avoid false negatives and extremely low positive counts, we considered compounds annotated with multiple MOAs or multiple gene targets as a positive match if one of their annotations overlapped with compounds with singleton annotations.

We calculated average precision by comparing Pearson correlation to ground truth annotations. Average precision calculates the mean of precision at each threshold in a precision-recall curve. We used the default “macro” average method in scikit-learn version 0.24.2 to calculate average precision, which does not weight precision means per label. We repeated this procedure without the same-dose constraint, designating repli

#### Supervised mechanism of action prediction: Multilabel-classification framework

We structured the classification task to predict compound MOAs and GO terms from different input profiling modalities. Specifically, we created a binary label matrix for each individual MOA or GO term with corresponding labels for each compound. This formulation created a multi-label framework because many compounds have previously been annotated with two or more specific mechanisms (Corsello et al., 2017). For example, if a compound is annotated to mechanism “A” and mechanism “B”, the binary matrix would include positive labels for two different columns.

We used Cell Painting and L1000 profiles to predict the same MOA or GO term binary matrix. In all cases, we used level 4 replicate profiles as input for model classification. For Cell Painting, we used feature-selected spherized profiles (level 4bs) and for L1000 we used non-spherized profiles. We treated each input datasets in exactly the same fashion as we describe in the subsections below.

#### Supervised mechanism of action prediction: Training and test splits

In order to prevent signal leakage from the training set into the test set, we carefully split the compounds as input into the training and test sets. Specifically, we first split compounds based on MOA count. This means, for example, that if an MOA was represented by just one compound, we placed that compound in the training set. However, if an MOA had more than one compound, we split the compounds for that individual MOA between training and test set based on the 80/20 train/test ratio. Because some compounds are annotated to more than one MOA (hence “multi-label”), we needed to iterate, repeating the random splits, until these conditions were satisfied for all MOAs. Ultimately, this results in zero overlap of compounds in the training set compared to the test set. We used the same exact training and test set compounds for each assay.

To ascertain and verify that the classification models are learning from the training set and that they could generalize well on test set data, we created a shuffle data set using data in the training set. The shuffle data set consists of the same features and data as the normal training set, but we randomly shuffled target labels. We provided incorrect MOAs for all replicate profiles, and retrained and reevaluated all models on the same tasks.

#### Supervised mechanism of action prediction: Cross validation and model selection

To account for class imbalance in compound replicates in each multi-label MOA in the training set, we divided the compounds into two major groups based on treatment replicate count: less frequent and highly frequent. The same compound may be annotated to multiple MOAs, but we considered each MOA label independently when splitting data for cross validation.

We then applied a 5-fold double-stratified cross-validation strategy to the training set. We split compounds across cross-validation folds balanced by MOA (or GO term) and according to compound replicate count. Specifically, we assigned replicates of less frequent compounds to the same fold, but we distributed replicates of highly frequent compounds evenly across folds. The threshold for dividing the compounds into less frequent and highly frequent categories is 20 for L1000 and 25 for Cell Painting. This threshold number means if the compound is found in less than 20 or 25 replicates in the training set it is considered less frequent, otherwise it is considered highly frequent. In practice, most compounds in our training set belonged to the “less frequent” category. This procedure, termed “drug stratification” in the MOA Kaggle competition (Kaggle.com et al., 2020), caused our training folds to be evenly distributed by MOA category and to mostly contain unique compound perturbations, which encourages models that generalize to never-before-seen compounds. We used our cross validation strategy to select optimal hyperparameters, but only evaluated models using a held-out test set of unique compounds.

#### Supervised mechanism of action prediction: Model architecture

We chose models for the multi-label MOA predictions from the top-2 winners from the MOA Kaggle competition (Kaggle.com et al., 2020). The models included 1D-Convolutional Neural Network (1D-CNN), TabNet (Attentive Interpretable Tabular Learning), Residual Neural Network (ResNet) and Simple Neural Network (Simple-NN) (Arik and Pfister, 2019; Fukushima, 1980; He et al., 2016; LeCun et al., 2015). We modified the winning architectures to handle different assay input dimensions.

Specifically, the 1D-CNN architecture consisted of four convolutional layers with kernel sizes of 3 and 5, stride of 1 and padding sizes of 2 and 1. We added adaptive and max pooling layers, as well as batch normalization (Ioffe and Szegedy, 2015) and drop-out layers within the convolutional architecture to encourage better model generalization. The TabNet architecture consisted of a width of 64 for the decision prediction layer, a width of 128 for the attention embedding for each mask, 1 step in the architecture and gamma value of 1.3. The ResNet architecture consisted of six fully-connected layers with batch normalization and drop-out layers included within the architecture. We used rectified linear units (RELU) and exponential linear units (ELU) as activation functions between layers (Agarap, 2018; Clevert et al., 2015). The Simple-NN architecture consisted of three fully-connected layers accompanied with batch normalization layers, drop-out and linear activation function layers. The optimization phase for all the models was done using Adam Optimizer with varying learning rates (Kingma and Ba, 2014). We independently optimized each architecture using data from each assay using the cross validation strategy as described above.

We also used an ensemble of the above-mentioned models in the MOA predictions by combining individual model predictions (weighted equally), then averaging the predictions to get an ensemble/blended version of all the models. We used multi-label k-nearest neighbors (K-NN) as a baseline model to compare performance (Altman, 1992; Fix and Hodges, 1951).

For complete details of all architectures and implementation instructions, refer to https://github.com/broadinstitute/lincs-profiling-complementarity (Way et al., 2021a).

#### Supervised mechanism of action prediction: Feature engineering and data normalization

Prior to model training, we added features to the training and test sets. These features included principal components, UMAP features, factor analysis components, and statistical features such as sum, mean, kurtosis and standard deviation of all the features, for all four input datasets. Specifically, we added 25 UMAP features and 50 factor analysis components from the existing data prior to the Simple-NN, and we added 25 principal components to the 1D-Convolutional Neural Network, TabNet and ResNet models. Lastly, we added statistical features to the TabNet model. We normalized all features using z-score normalization prior to model training.

#### Supervised mechanism of action prediction: Model evaluation

The output of all the models is a probabilistic value between 0 and 1 corresponding to the probability of the model predicting a given MOA class label. We evaluated models calculating area under the Precision-Recall curve (AUPR). AUPR is a threshold-invariant metric that takes into account recall and precision, of which precision is particularly important because it measures the fraction of correct predictions among the positive predictions. AUPR accounts for imbalanced datasets, which is useful for evaluating classification tasks in highly imbalanced datasets (Saito and Rehmsmeier, 2015). We also calculated AUPR in randomly shuffled MOA class labels. To create this randomly shuffled matrix, we kept the MOA label count the same per MOA. To prevent unbalanced evaluation metrics in the test set, we removed bortezomib (positive control) from all evaluations.

We used micro-averaging in our AUPR calculation for both “global” performance and per-MOA metrics. For the “global” AUPR (total performance) we aggregate the contributions of all compounds to compute the average metric. For the per-MOA AUPR we aggregate the contributions of all compounds annotated to the specific MOA.

#### Supervised Gene Ontology term prediction

For predicting GO terms, we repeated the same supervised learning procedures as described above for compound MOAs. A major step that was different for the GO term analysis was the requirement to map compound target annotations to GO terms.

In addition to MOA annotations, The Drug Repurposing Hub also includes gene target annotations for most compounds. We mapped these gene target annotations to all GO Biological Process, GO Molecular Function, and GO Cellular Component terms (GO.db version 3.14.0 (Carlson, 2017a)) using topGO version 2.46.0 (Adrian Alexa, 2017; Alexa et al., 2006) and org.Hs.eg.db version 3.14.0 (Carlson, 2017b). The same compound may be annotated to multiple gene targets, and we considered each gene target label independently. For example, if a compound targeted HDAC1 and PIK3CA, we considered that compound to belong to all GO terms containing both genes. We only considered GO terms for downstream supervised learning predictions if the term had 20 or more unique compound annotations. This procedure resulted in a total of 773 GO terms to predict.

## QUANTIFICATION AND STATISTICAL ANALYSIS

We performed all statistical analysis using the packages outlined in the Key Resources table and can be reproduced in full using the details outlined in the computational reproducibility and data availability sections. We chronologically discuss complete descriptions of all statistical procedures in relevant results and/or methods sections.

## Supplementary Tables

Table S1: Perturbation identifiers with mechanism of action

Table S2: Perturbation reproducibility across assays and doses

Table S3: Gene overrepresentation analyses of perturbations that consistently alter cell morphology

Table S4: Metrics for matching compounds based on mechanism of action

Table S5: Metrics for matching compounds based on gene targets

Table S6: Deep learning performance for predicting perturbation mechanism of action

