## Supplementary Figures for "Morphology and gene expression profiling provide complementary information for mapping cell state"

**Figure S1.** *Cell Painting example images, related to Figure 1.* **3**

**Figure S2.** *L1000 example probe expression, related to Figure 1.* **4**

**Figure S3.** *Data collection and data processing workflows, related to Figure 1.* **5**

**Figure S4.** *Comparing cell count and replicate reproducibility in Cell Painting data, related to Figure 1.* **6**

**Figure S5.** *Plate position effects are reduced by a spherize transform in Cell Painting, related to Figure 1.* **7**

**Figure S6.** *Walking through the percent replicating metric, related to Figure 1.* **8**

**Figure S7.** *Percent replicating scores with different input Cell Painting and L1000 data, related to Figure 1.* **9**

**Figure S8.** *Percent strong scores with different input Cell Painting and L1000 data calculated using different null distributions, related to Figure 1.* **10**

**Figure S9.** *Plate map non-replicate diffusion sampling indicates minor plate position effects, related to Figure 1.* **11**

**Figure S10.** *Cell Painting UMAP of select compound mechanisms of action (MOA), related to Figure 2.* **12**

**Figure S11.** *L1000 UMAP of select compound mechanisms of action (MOA), related to Figure 2.* **13**

**Figure S12.** *Alternative lower dimensional embeddings for Cell Painting and L1000 profiles, related to Figure 2.* **14**

**Figure S13.** *Principal components analysis (PCA) identifies spaces associated with compound reproducibility, related to Figure 2.* **15**

**Figure S14.** *UMAP for a second batch of Cell Painting profiles including alternate timepoints and cell lines, related to Figure 2.* **16**

**Figure S15.** *Goodness of fit clustering metrics for Cell Painting and L1000, related to Figure 3.* **17**

**Figure S16.** *Tracking feature variation in Cell Painting and L1000 profiles, related to Figure 3.* **18**

**Figure S17.** *Cell Painting and L1000 differentially measure compound perturbations by mechanism of action (MOA), related to Figure 4.* **19**

**Figure S18.** *Poor performances for mechanism of action prediction, related to Figure 5.* **20**

**Figure S19.** *Predicting compound gene targets annotated to Gene Ontology (GO) terms in Cell Painting and L1000 reveals overlapping and complementary performance for different pathways, related to Figure 5.* **21**

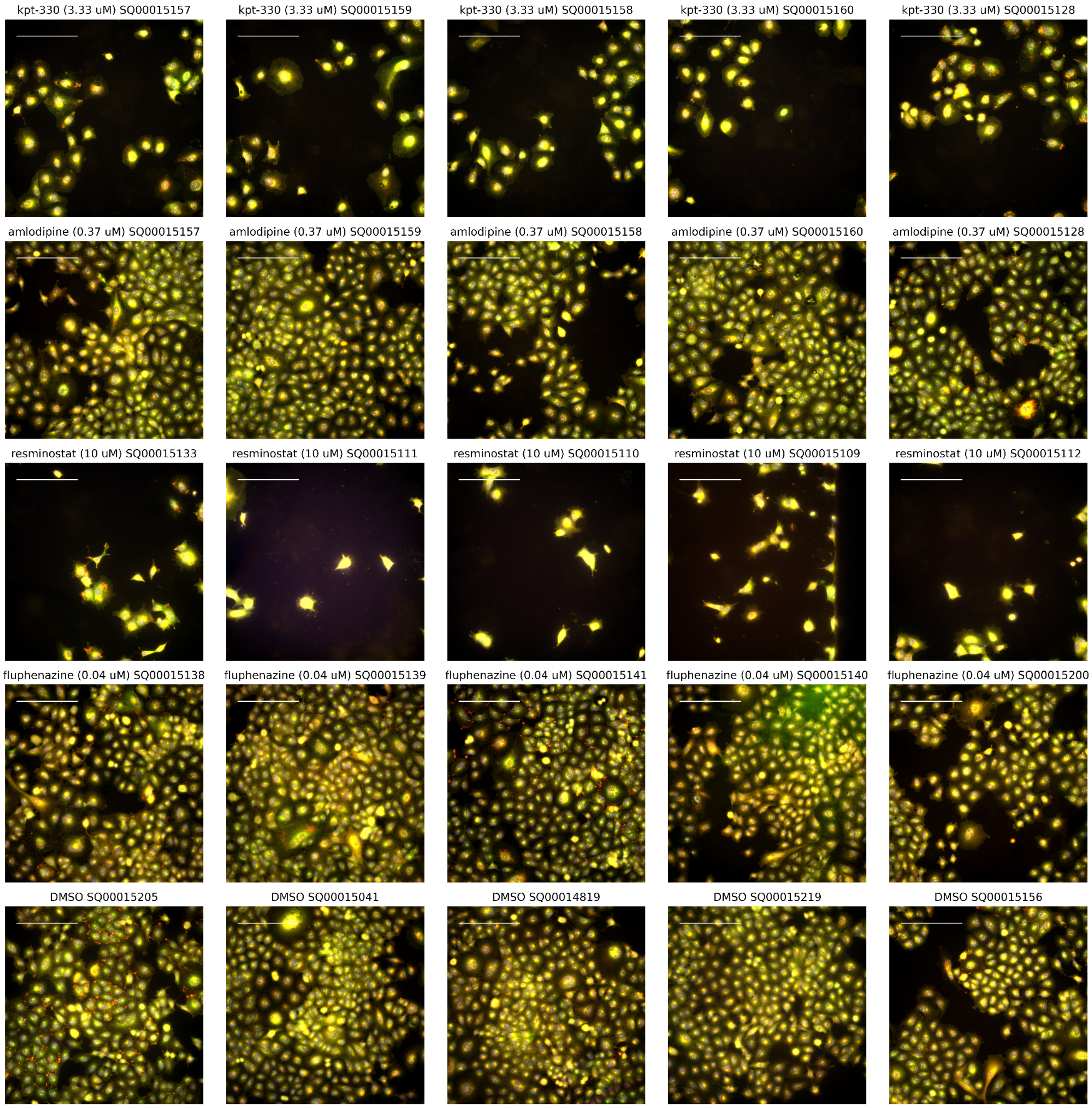


**Figure S1.** *Cell Painting example images, related to Figure 1.*

We provide example images for four representative compound perturbations and the negative control DMSO. For compounds, we selected the most reproducible treatment in Cell Painting (top row: KPT-330 3.33 uM), the most reproducible treatment in L1000 (second row; Resminostat 10uM), the least reproducible treatment in Cell Painting (third row; amlodipine 0.37uM), and the least reproducible treatment in L1000 (fourth row; Fluphenazine 0.04uM). We randomly selected different plates and wells and showed the most central field of view (FOV) in each random well. We provide merged representations of the five Cell Painting channels (see Figure 1). Scale bar in the top left is 200 µm.

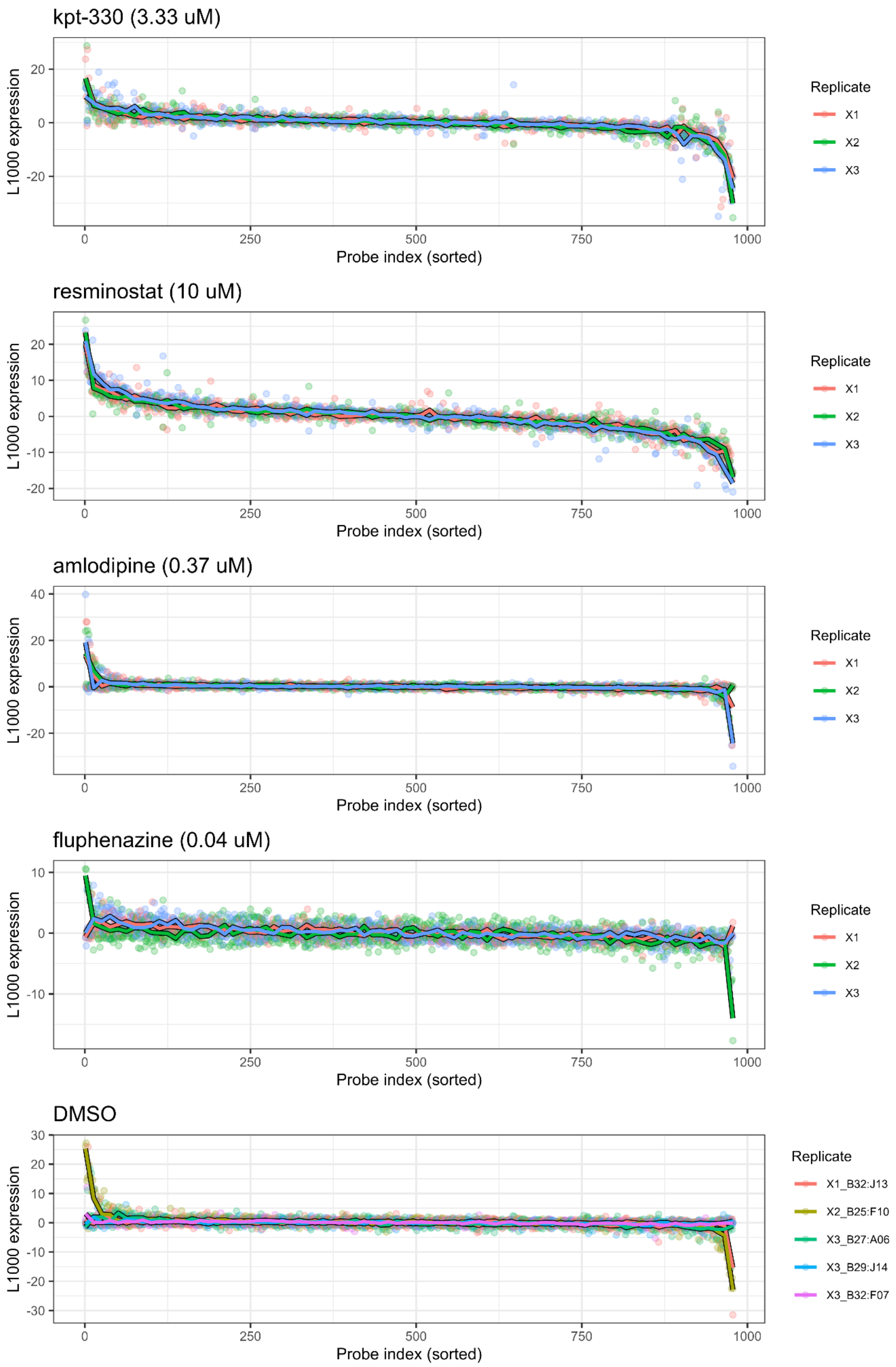


**Figure S2.** *L1000 example probe expression, related to Figure 1.*

We provide example expression values across 978 landmark genes of the L1000 assay for four representative compound perturbations and the negative control DMSO. For compounds, we selected the most reproducible treatment in Cell Painting (top row: KPT-330 3.33 uM), the most reproducible treatment in L1000 (second row; Resminostat 10uM), the least reproducible treatment in Cell Painting (third row; amlodipine 0.37uM), and the least reproducible treatment in L1000 (fourth row; Fluphenazine 0.04uM). We randomly selected five DMSO samples to display. We sorted the probe index by total expression values independently per row.

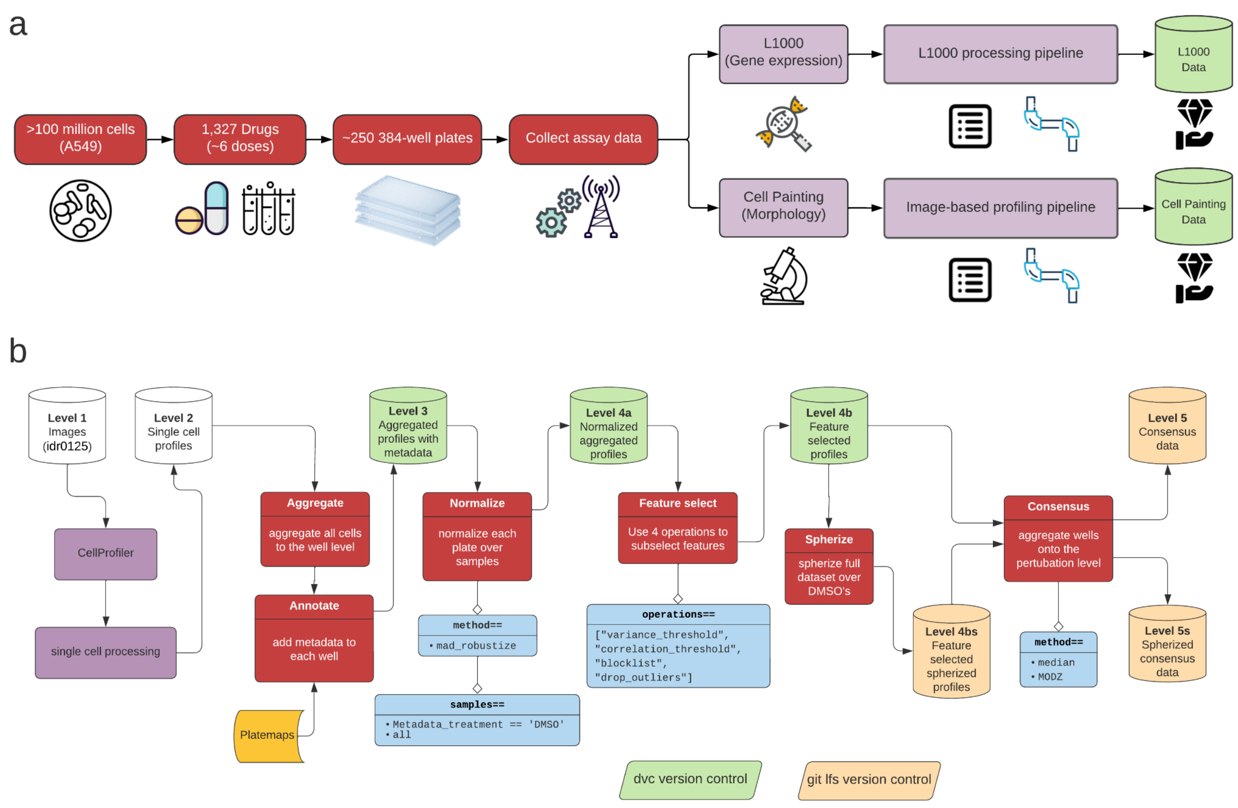


**Figure S3.** *Data collection and data processing workflows, related to Figure 1.*

**(a)** We cultured A549 lung cancer cells and exposed them to 1,327 different compound perturbations in about six doses per compound. We plated these cells in 384 well plates, and, using the same plate layout, measured gene expression (using the L1000 assay) and morphology (using the Cell Painting assay) in compound-perturbed A549 cells. **(b)** Our image-based profiling pipeline we used to process the Cell Painting images. We used pycytominer to process the single cell profiles. All processing code and profile data are available at https://github.com/broadinstitute/lincs-cell-painting. Image data available at Image Data Resource (accession: idr0125).

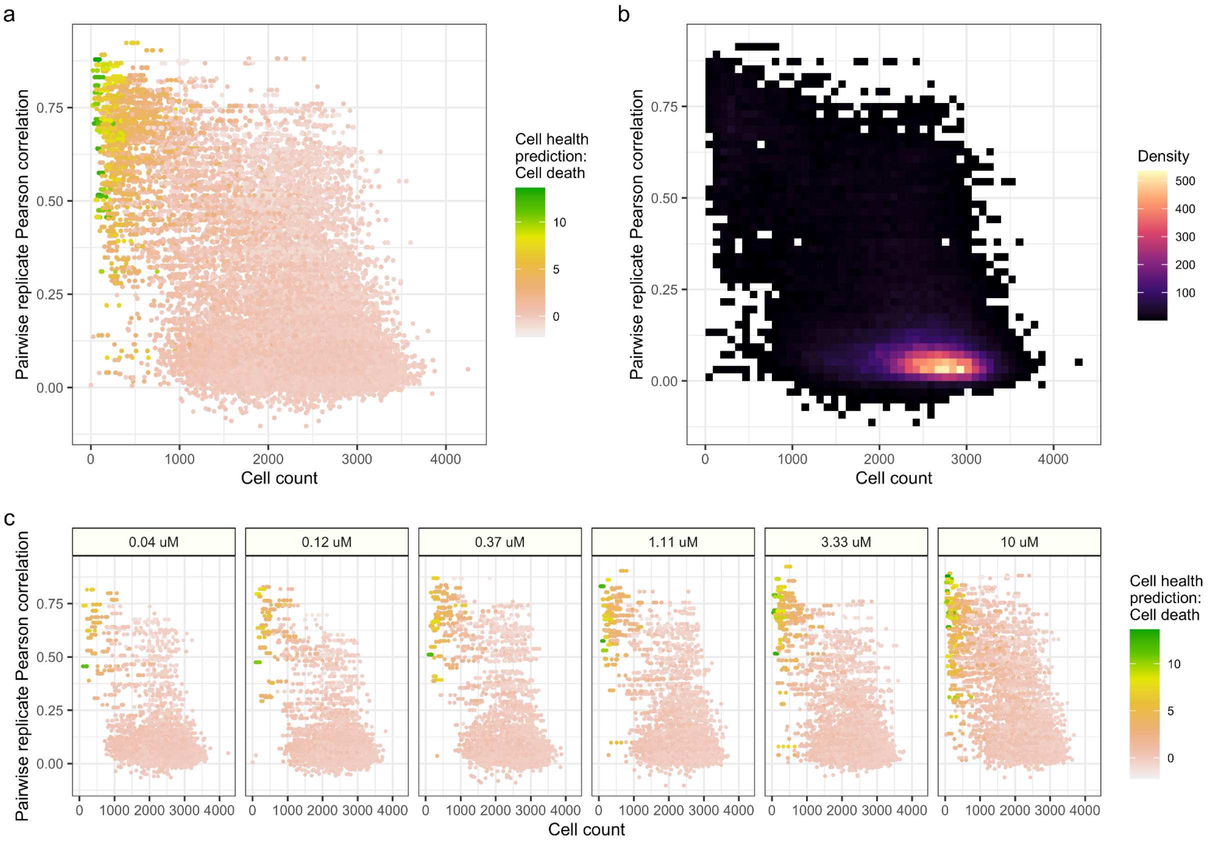


**Figure S4.** *Comparing cell count and replicate reproducibility in Cell Painting data, related to Figure 1.*

**(a)** All profile summary. **(b)** The majority of profiles have between 2,000 and 3,000 cells and low replicate reproducibility. **(c)** Comparing cell count and reproducibility across dose. We used previously validated cell death prediction models in panels a and c. We used machine learning predictions from the Cell Death machine learning model from Way et al. 2021.

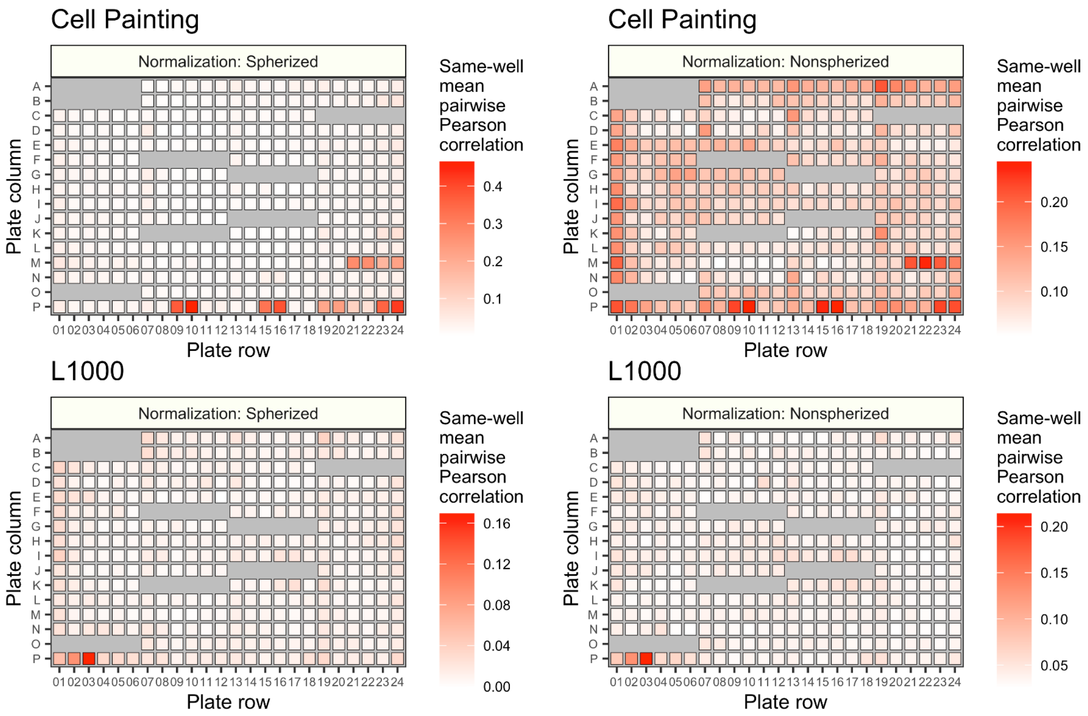


**Figure S5.** *Plate position effects are reduced by a spherize transform in Cell Painting, related to Figure 1.*

We calculated mean pairwise correlations of non-replicate (not-same-compound) profiles that were collected from the same well, but a different plate map. Wells with missing boxes indicate wells that always contained replicates of the same treatment, across all plate maps (controls). We observed same-well position effects throughout non-spherized Cell Painting data (top right). After applying a spherize transform, we observed a reduction, but not complete ablation, of plate position artifacts (top left). We did not observe much of an impact of spherizing the data for L1000 profiles (bottom). Note the different color scales for each plate map figure.

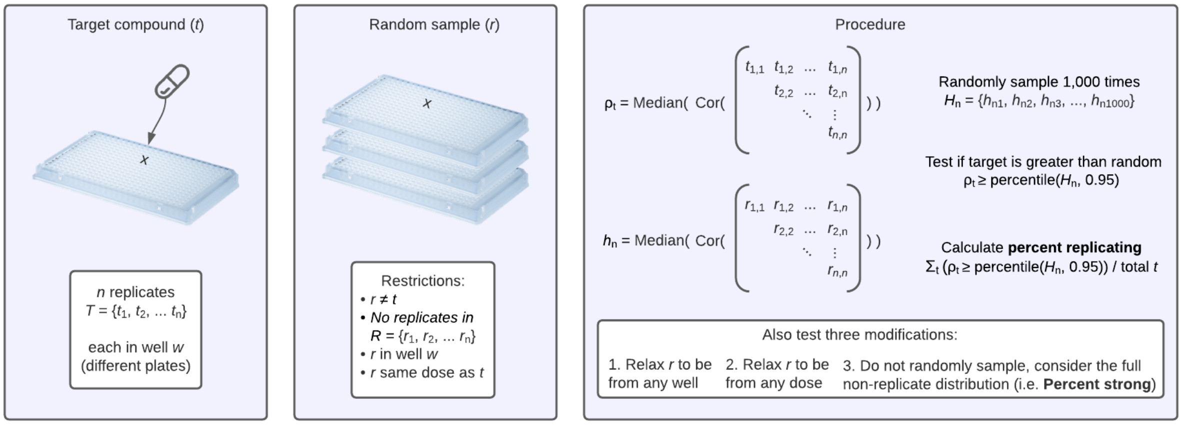


**Figure S6.** *Walking through the percent replicating metric, related to Figure 1.*

Percent replicating is a summary statistic per assay, which reports the percentage of target compounds whose replicates had higher pairwise correlations than random. The metric compares the median pairwise Spearman correlation of target compound replicates to 1,000 randomly sampled non-replicates with various sampling restrictions. We modify percent replicating in three key ways, and report how the metric behaves in these different scenarios (see **Figure S7**).

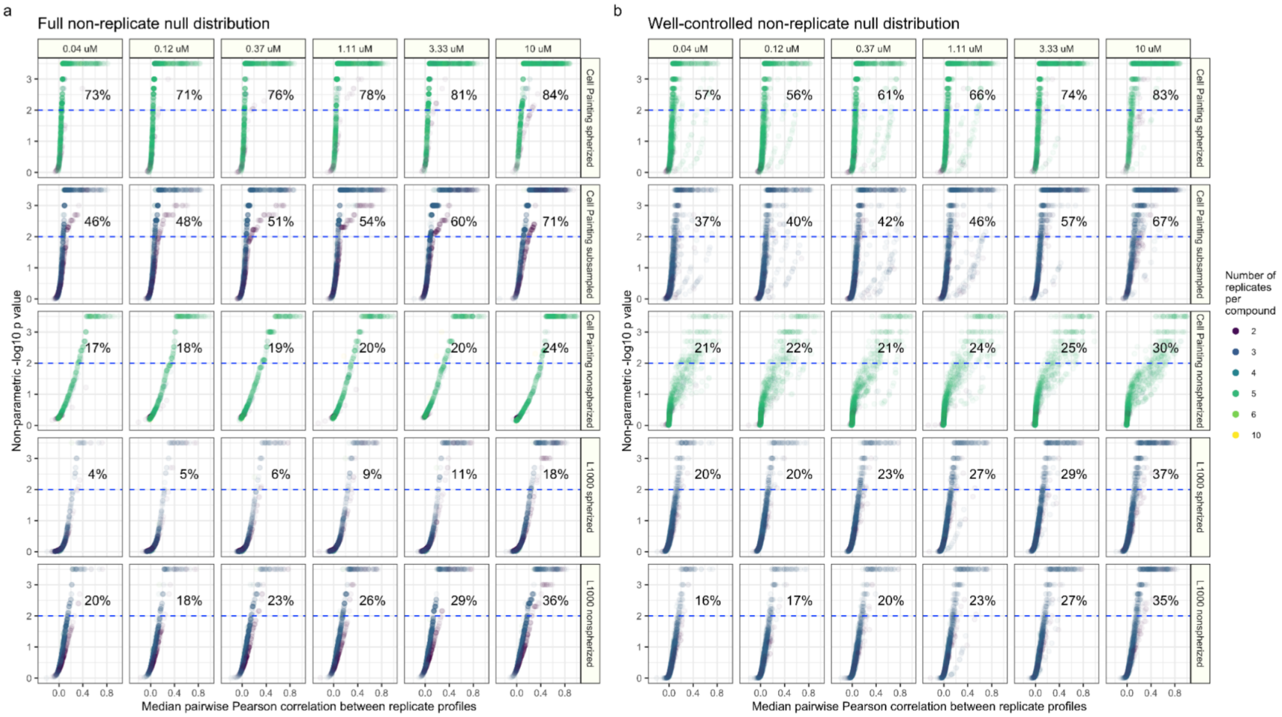


**Figure S7.** *Percent replicating scores with different input Cell Painting and L1000 data, related to Figure 1.*

We calculated percent replicating scores for five different input data sets (rows, top to bottom): (1) All Cell Painting replicates with spherize transform (same data as figure 1A), (2) Randomly subsampled Cell Painting replicates to match the number of replicates per treatment (compound by dose) in the L1000 assay, (3) All Cell Painting replicates without spherize transform, (4) All L1000 replicates with a spherize transform, and (5) All L1000 replicates without a spherize transform (same data as figure 1A). We calculated percent replicating using two carefully matched null distributions (see methods) per input dataset. **(a)** Using a replicate cardinality-matched median non-replicate pairwise correlation null distribution. **(b)** Using a replicate cardinality and same-well matched median non-replicate pairwise correlation null distribution.

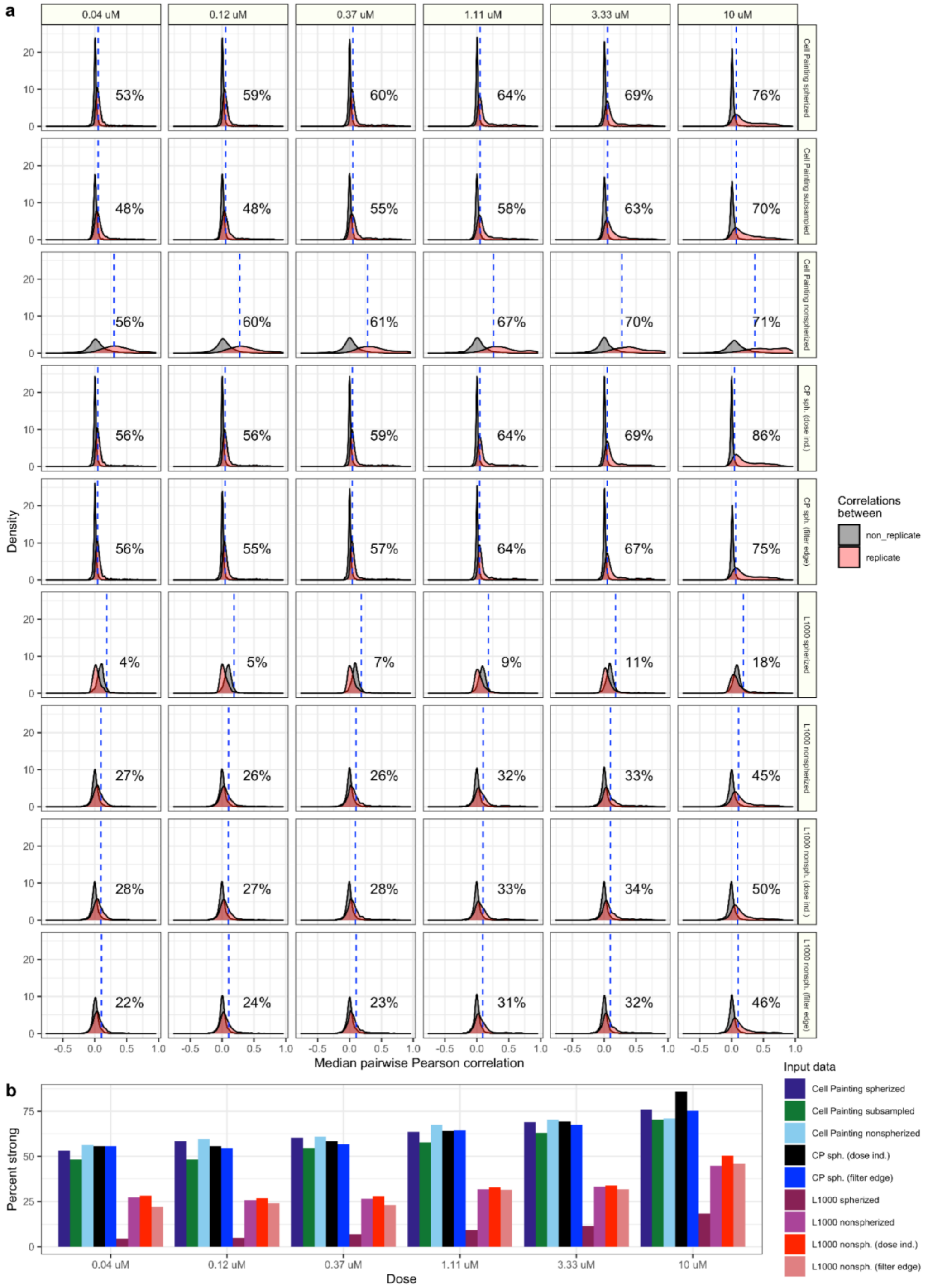


**Figure S8.** *Percent strong scores with different input Cell Painting and L1000 data calculated using different null distributions, related to Figure 1.*

We calculated percent strong (see Methods) for profiling readouts across the data sets. **(a)** Replicate and non-replicate distributions of median pairwise correlations of the nine inputs as specified above constructed with various null distribution sampling strategies. **(b)** Percent strong summary.

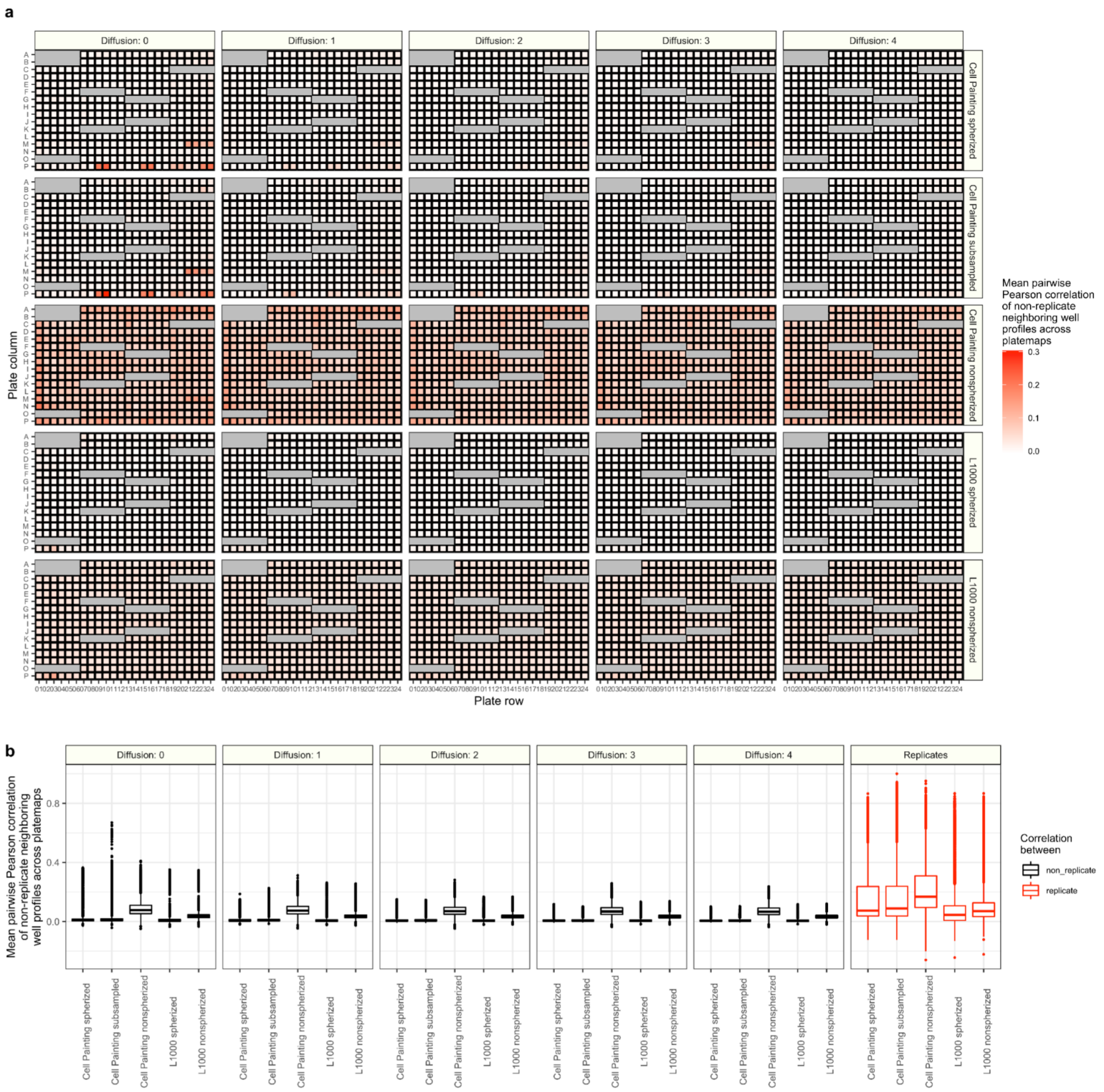


**Figure S9.** *Plate map non-replicate diffusion sampling indicates minor plate position effects, related to Figure 1.*

In a diffusion analysis, we systematically increased the local neighborhood of wells to measure mean non-replicate Pearson correlations across different plate maps. **(a)** Increasing the local neighborhood slightly decreases mean non-replicate pairwise Pearson correlations with some bias towards edge wells. A spherize transform dramatically reduces this non-replicate correlation. **(b)** A summary of all wells across diffusion parameters for replicates (red) and non-replicates (black). In general, the replicate correlations are substantially higher than non-replicate correlations.

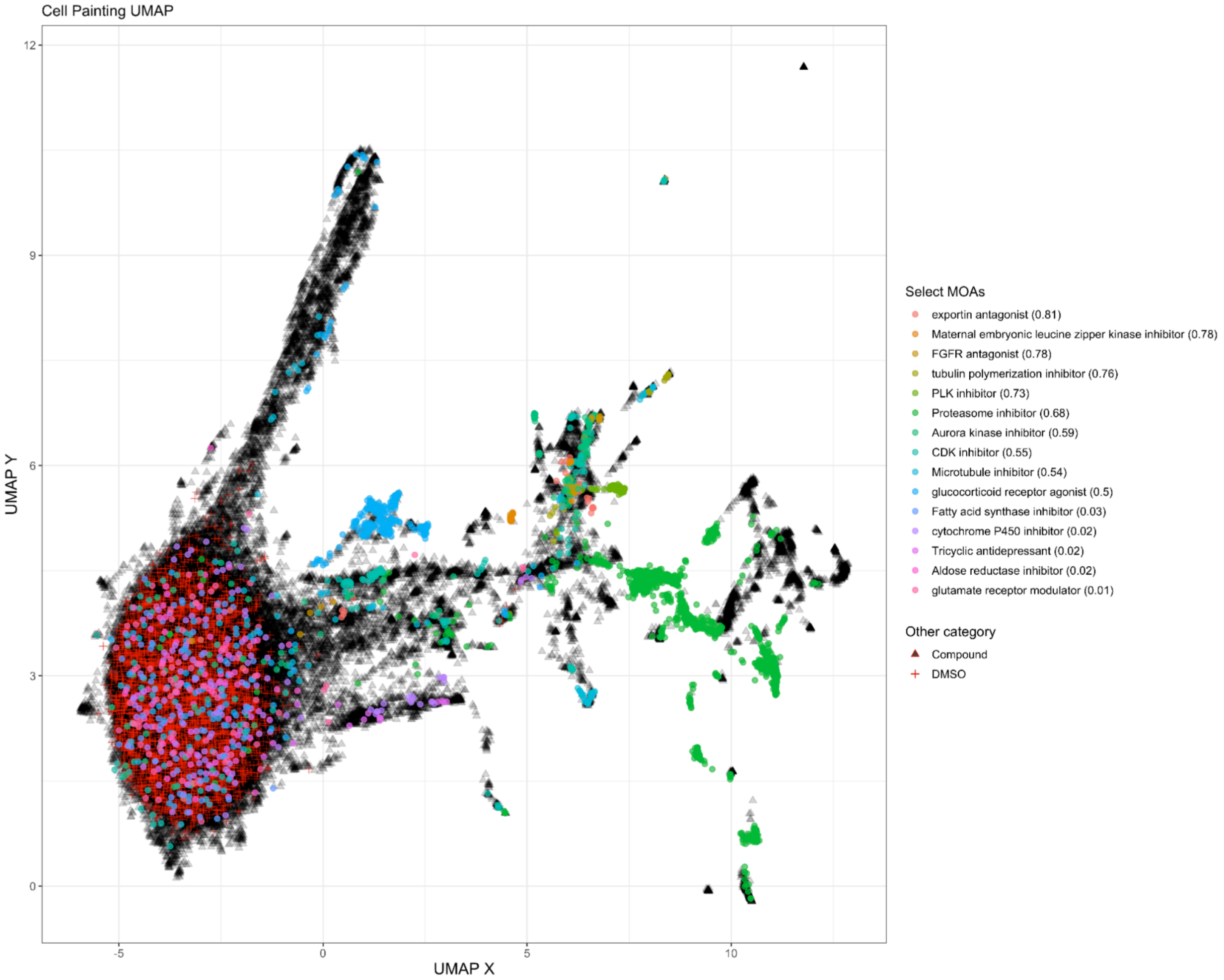


**Figure S10.** *Cell Painting UMAP of select compound mechanisms of action (MOA), related to Figure 2.*

Highlighting all compounds annotated with the top 10 and bottom 5 reproducible MOAs based on compound replicate pairwise correlation. We used specific filtering criteria to select which compounds to visualize. 1) Select the top 5 MOAs regardless of sample size; 2) Select the next top 5 MOAs (without replacement) but each MOA needed to have at least 30 compound representatives; 3) Select the least correlating MOAs with at least 30 compound representatives. The number next to each MOA in the legend represents the median value of all annotated compounds' median pairwise correlations. The UMAP depicts level 4 replicate Cell Painting profiles across all doses.

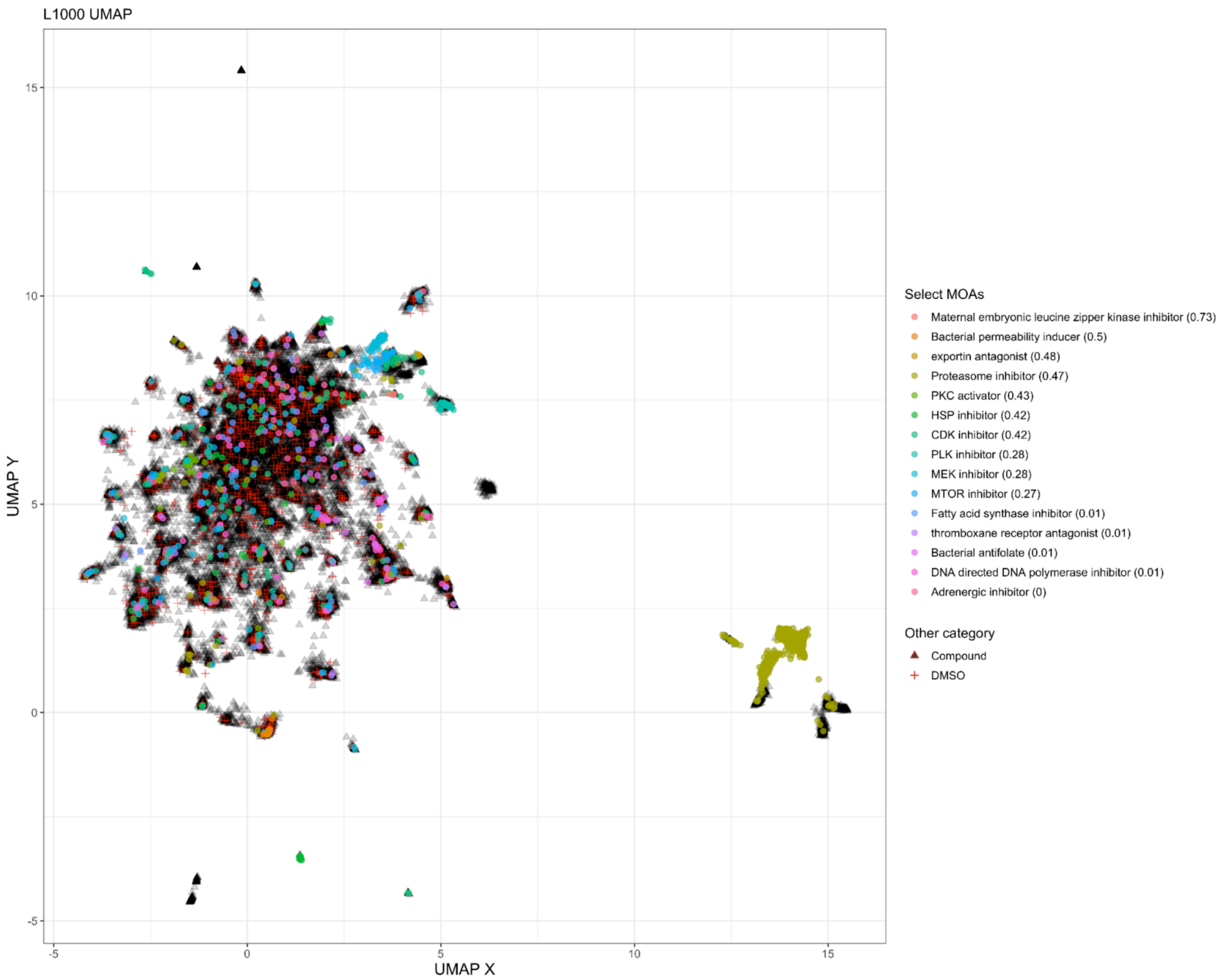


**Figure S11.** *L1000 UMAP of select compound mechanisms of action (MOA), related to Figure 2.*

Highlighting all compounds annotated with the top 10 and bottom 5 reproducible MOAs based on compound replicate pairwise correlation. We used specific filtering criteria to select which compounds to visualize. 1) Select the top 5 MOAs regardless of sample size; 2) Select the next top 5 MOAs (without replacement) but each MOA needed to have at least 18 compound representatives; 3) Select the least correlating MOAs with at least 18 compound representatives. The number next to each MOA in the legend represents the median value of all annotated compounds' median pairwise correlations. The UMAP depicts level 4 replicate L1000 profiles across all doses.

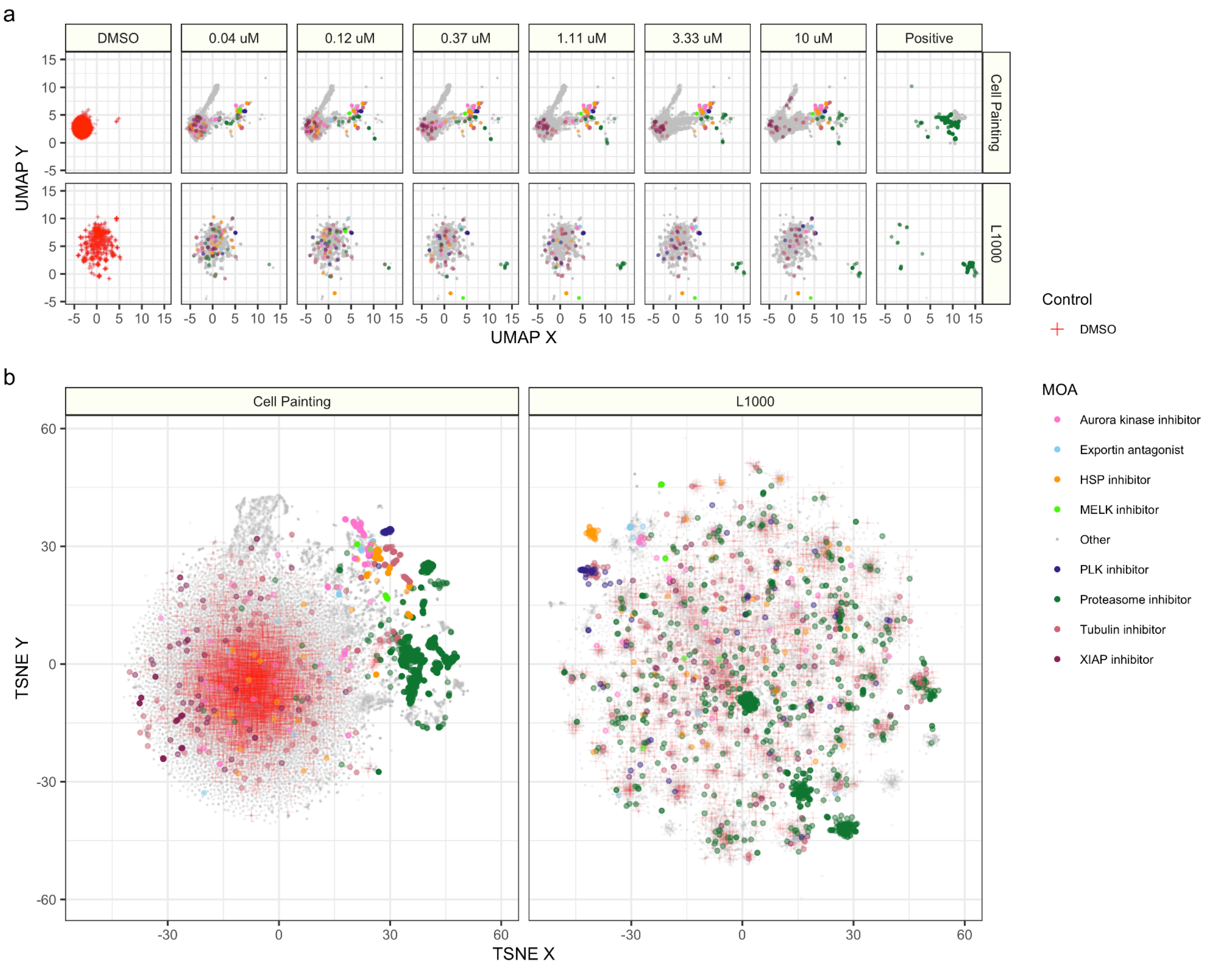


**Figure S12.** *Alternative lower dimensional embeddings for Cell Painting and L1000 profiles, related to Figure 2.*

We fit a Uniform Manifold Approximation (UMAP) and t-distributed stochastic neighbor embedding (TSNE) on Cell Painting and L1000 data (level 4 replicates) separately using the full data. **(a)** UMAP coordinates by dose. **(b)** TSNE coordinates across all doses. We show DMSO and positive controls separately from dose to better highlight patterns from non-control compounds.

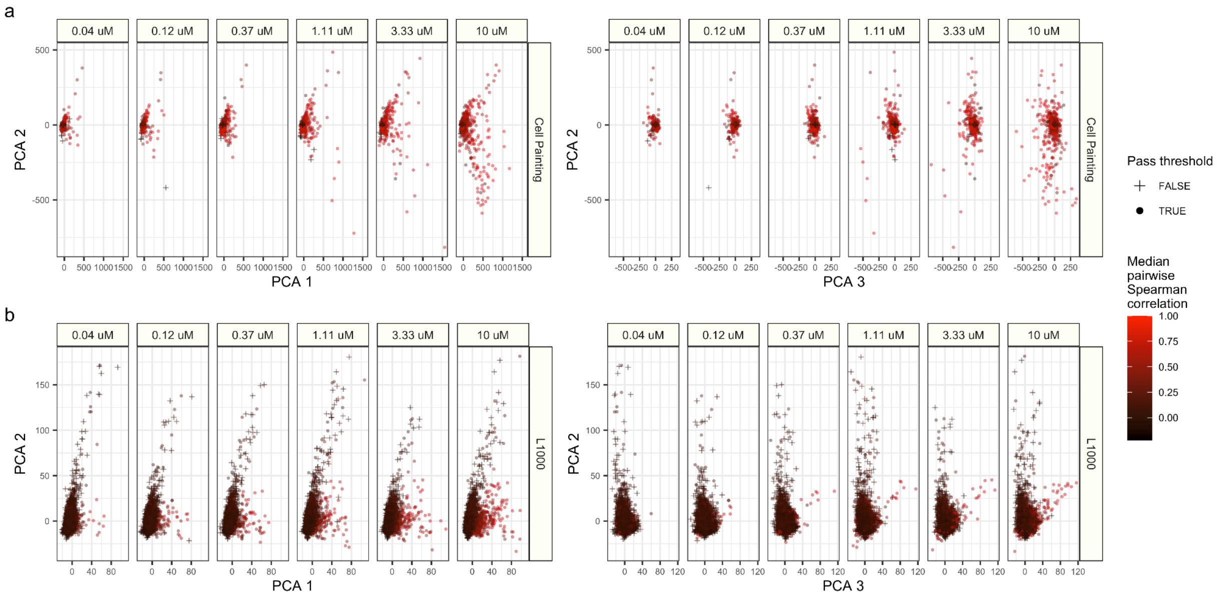


**Figure S13.** *Principal components analysis (PCA) identifies spaces associated with compound reproducibility, related to Figure 2.*

The top three components for the **(a)** Cell Painting and **(b)** L1000 assays and corresponding median pairwise Spearman correlations per treatment.

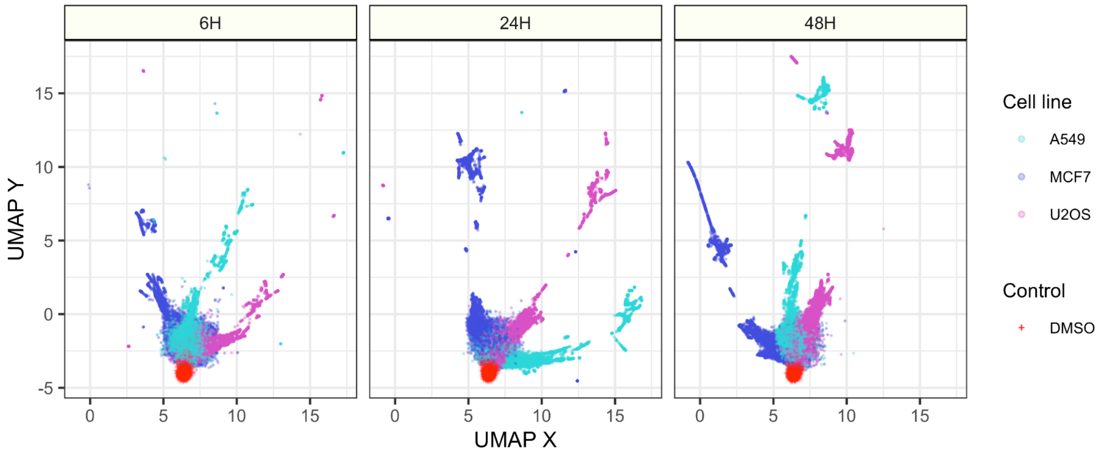


**Figure S14.** *UMAP for a second batch of Cell Painting profiles including alternate timepoints and cell lines, related to Figure 2.*

We fit a Uniform Manifold Approximation (UMAP) on batch 2 of level 4 Cell Painting profiles. These data were treated with three different incubation periods (6 hours, 24 hours, and 48 hours) and included three different cell lines. The tendency of all conditions of each cell line to block together shows that differences between cell lines typically outweigh those caused by treatments. We spherized these profiles independently by incubation time and cell line. We indicate all DMSO controls for all three cell lines in red (they are all clustered similarly).

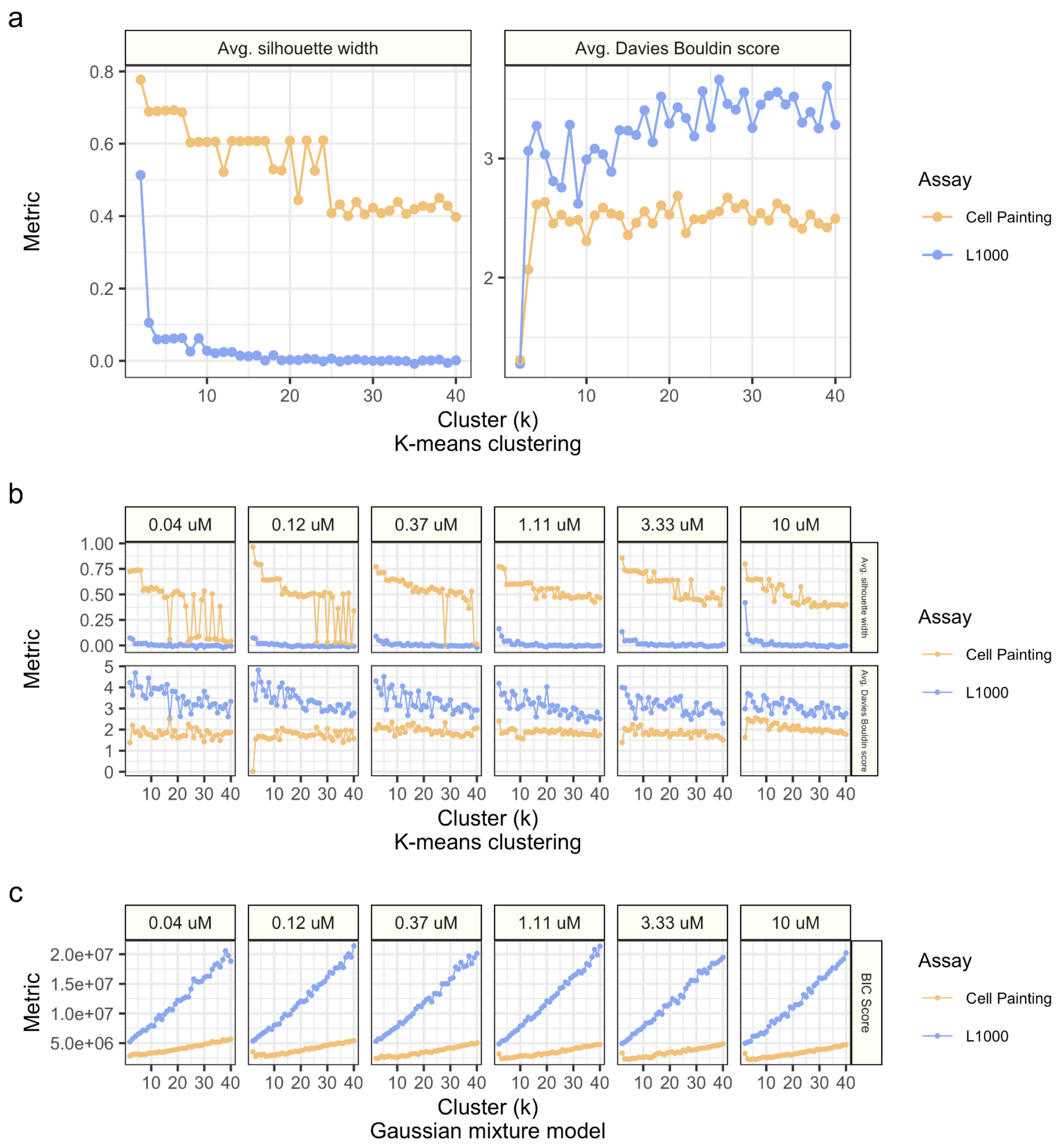


**Figure S15.** *Goodness of fit clustering metrics for Cell Painting and L1000, related to Figure 3.*

Using the 1,327 common compounds (level 4 replicate profiles), we fit clustering algorithms to identify k=2 through k=40 clusters. Prior to clustering, we transformed both input data using principal component analysis (PCA) using 350 PC dimensions. Using k-means clustering, we report the average Silhouette width and average Davies Bouldin scores for clustering solutions using compound treatments from **(a)** all doses and **(b)** each dose independently. Higher silhouette scores indicate better separation between clusters while higher Davies Bouldin scores indicate higher similarity between clusters. **(c)** Fitting Gaussian mixture models (GMM), we acquire Bayesian information criterion (BIC) scores to report on clustering goodness-of-fit (see methods; lower is better) for clustering solutions with k=2 through k=40. We indicate the specific metrics in panels (b) and (c) in each right facet label.

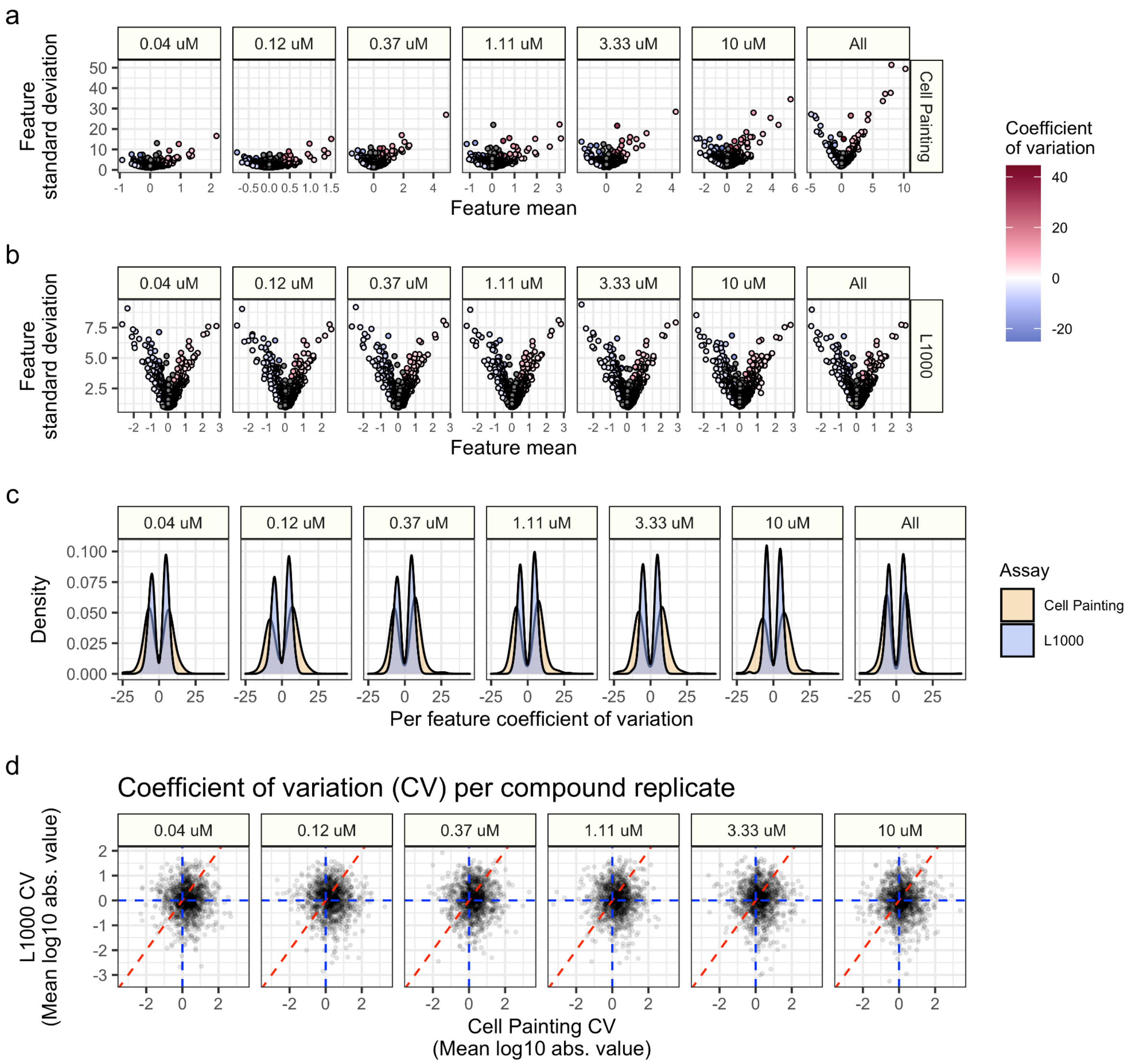


**Figure S16.** *Tracking feature variation in Cell Painting and L1000 profiles, related to Figure 3.*

Feature mean, standard deviation, and coefficient of variation (CV; standard deviation / mean) in **(a)** Cell Painting and **(b)** L1000 profiles. Note the different axis scales and common legend colors. Each point represents a feature. Features with a mean between -0.3 and 0.3 are grayed out in order to maintain an interpretable color scale (removing color from features with extremely large CV values). **(c)** CV distributions for Cell Painting and L1000 profiles demonstrate higher relative variability in Cell Painting features. The bimodality reflects the fact that CV takes the sign of the mean of the feature values, which end up being about equally split between negative and positive. **(d)** Per replicate CV comparing replicate treatments for Cell Painting (x axis) and L1000 (y axis). Each point represents the log10 mean absolute value CV for all features measured within each replicate treatment. We remove treatments with CV values equal to zero. All panels display level 4 Cell Painting spherized profiles and level 4 L1000 nonspherized profiles.

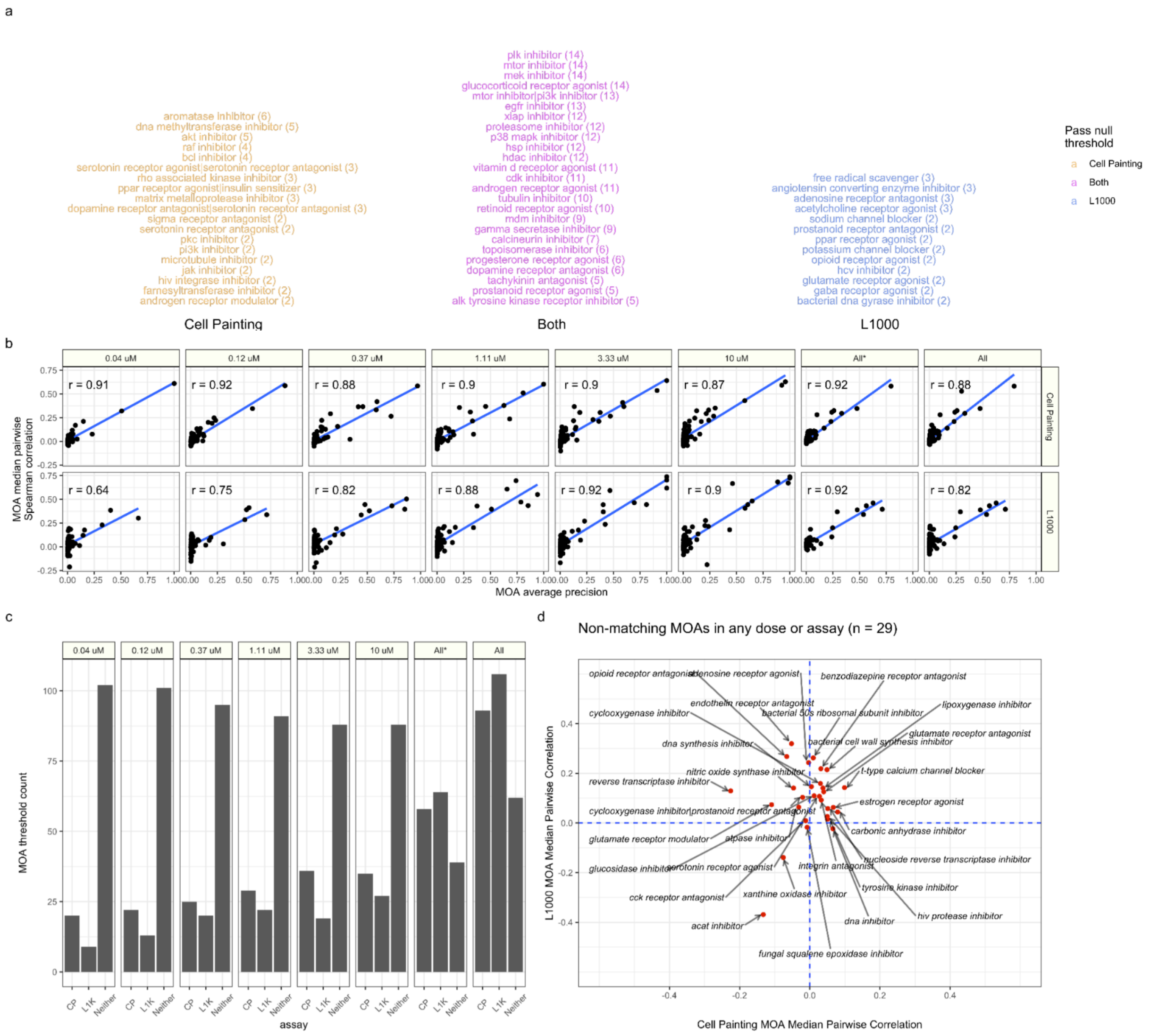

**Figure S17.** *Cell Painting and L1000 differentially measure compound perturbations by mechanism of action (MOA), related to Figure 4.*

**(a)** All MOAs that are consistently measured by Cell Painting, L1000, or both in at least two doses (including all dose comparison). Brackets indicate the number of times the MOA was reproducible (per dose and assay). For example, PLK inhibitors were reproducible in every dose and in each assay (6 doses + 1 all dose  x 2 assays = 14). **(b)** MOA median pairwise correlation and MOA average precision indicate similar high-performing MOAs. The All* bar represents matched MOAs for the 127 MOA set and the All bar represents matched MOAs for the 210 MOA set. **(c)** The number of MOAs that pass the percent matching null threshold for each dose. Within dose, the majority of MOAs do not match in either assay. Note, this is the same data shown in **Figure 4C**, with the number of MOAs that pass in "Neither" assay. **(d)** The 29 MOAs (out of 208) that are not reproducible in any dose or assay.

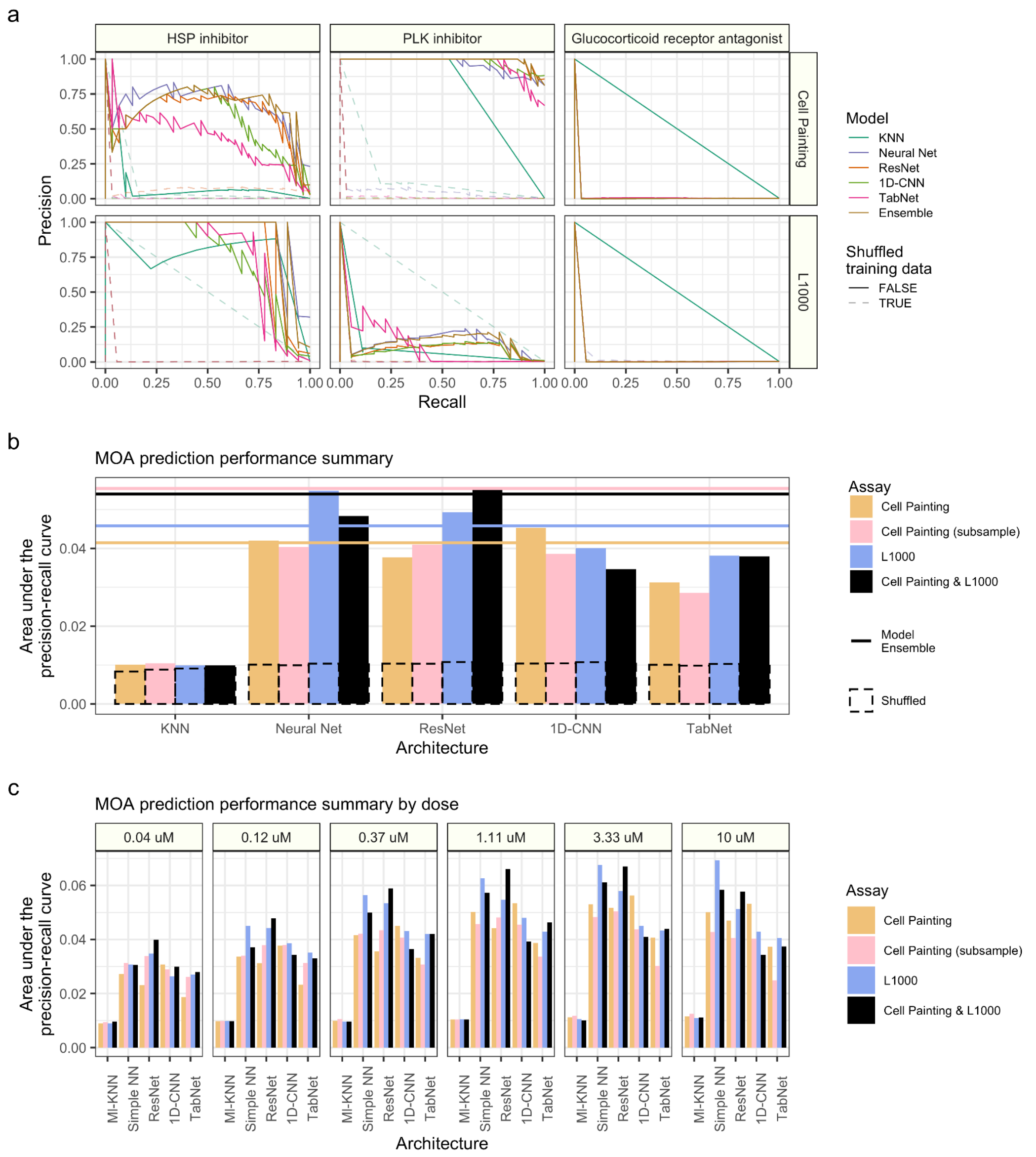


**Figure S18.** *Poor performances for mechanism of action prediction, related to Figure 5.*

**(a)** Precision-recall curve examples of poor performing mechanism of action (MOA) predictions in either or both assays. **(b)** Comparing individual assay performance to randomly subsampled Cell Painting data to match L1000 counts and both-assay concatenated model performance (see Methods). **(c)** We observed an increasing performance, by area under the precision recall curve (AUPR), with increasing dose.

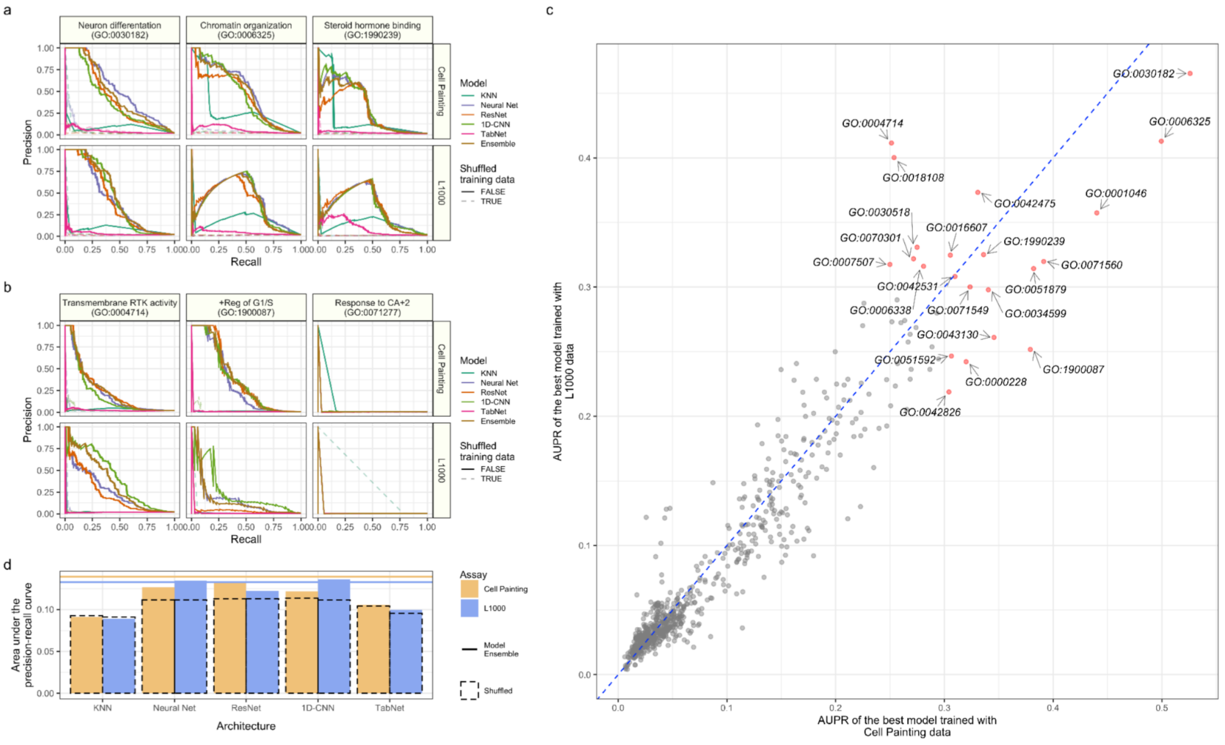


**Figure S19.** *Predicting compound gene targets annotated to Gene Ontology (GO) terms in Cell Painting and L1000 reveals overlapping and complementary performance for different pathways, related to Figure 5.*

**(a)** Held out test-set precision-recall curves for three well performing GO terms in both assays. **(b)** Held out test set precision-recall curve examples of poor performing GO term predictions in either or both assays. **(c)** Individual GO term performance by held out test set area under the precision-recall curve (AUPR) in the top performing model using Cell Painting and L1000 data. (**d**) Overall held out test set model performance measured by AUPR for GO term prediction for our multi-label, multi-class prediction framework. We trained models from a recent Kaggle competition plus a K nearest neighbors baseline model. The dotted bar chart represents a negative control in which we trained models with shuffled labels. The solid lines indicate ensemble model performance by blending model predictions (see Methods). We trained all models using level 4 replicate profiles.
